# Transcriptional responses to proteotoxic stressors are profoundly diverse and tissue-specific

**DOI:** 10.64898/2025.12.17.694942

**Authors:** Adelina Rabenius, Intisar Salim, Hilmar Lindström, Anastasiya Pak, Serhat Aktay, Anniina Vihervaara

## Abstract

Cells counteract proteotoxic conditions by launching transcriptional stress responses. While synthesis of Heat shock proteins (HSPs) upon acute stress is well-characterized, how distinct proteotoxic conditions reshape the transcriptome remains poorly understood. Here, we analyse polyA+ RNA expression under heat shock, HSP90 inhibition, and polyglutamine (polyQ) aggregation. We find fundamentally distinct transcriptional responses to proteotoxic stressors, and a systemic deficiency of mice under chronic stress to launch acute responses. While heat shock and HSP90 inhibition induce chaperones, polyQ aggregation increases RNAs linked to transcription repression, chromatin remodeling, and autophagy. Analysing wildtype and Huntington’s Disease (HD) mice reveals tissue-specific transcriptional adaptations to polyQ, including repressed cell-type specific functions and altered energy metabolism. Despite profound reprogramming, remarkably few RNAs are consistently induced (*Acy3*, *Abdh1*, *Tmc3*) or reduced (*Fos*) across HD brain regions. These results emphasize cellular background in disease manifestation, and support energy metabolism and detoxifying enzymes as therapeutic targets in late-stage HD. Moreover, the systemic deficiency of chronically stressed mice to launch responses challenges strategies that rely on induced transcription. Altogether, we characterize transcription signatures to proteotoxic stresses, identify key *trans*-activators driving proteotoxic stress responses, provide an interactive gene-by-gene viewer of global changes, and delineate tissue-specific transcription programs in HD mice.

## Introduction

Proteotoxic stress challenges homeostasis and, if unmitigated, leads to accumulation of misfolded and aggregated proteins. To counteract stress, cells launch protective responses, such as heat shock response (HSR) in the cytosol and nucleus, and unfolded protein response (UPR) in the endoplasmic reticulum and mitochondria (reviewed in Senft and Ronai, 2015). HSR is considered a rapid response to various stresses, including elevated temperatures, infections, heavy metals, protein aggregates, and chaperone incapacity (reviewed in Morimoto, 2008; Richter *et al*., 2010). However, the nature of distinct protein-damaging stresses can be profoundly different, requiring tailored responses. The most studied example, heat shock, triggers instant repression of transcription and translation, coupled to Heat shock factor 1 (HSF1) driven activation of chaperone, co-chaperone and polyubiquitin genes (reviewed in Vihervaara *et al*., 2018). The produced chaperone machineries, in turn, hold misfolded proteins, refold them into correct conformation, prevent aggregation, and direct misfolded proteins to proteasomal degradation or autophagy (reviewed in Kim *et al*., 2013).

Neurons are particularly susceptible to unbalanced proteostasis (Morimoto 2008), and thus, proteotoxic stress underlies neurodegenerative diseases such as Huntington’s Disease (HD). HD is a dominant genetic disease, inherited *via* abnormally long cytosine-adenine-guanine (CAG) repeats in *Huntingtin* (*Htt*) gene (MacDonald *et al*., 1993; reviewed in Walker, 2007). The length of the encoded glutamine repeats (polyQ) varies in the mutant protein (mHTT), the longer stretches causing an earlier onset and more severe symptoms (Walker, 2007). The underlying mechanisms of how mHTT drives the disease are not fully understood. However, HD patients display protein aggregates in the brain, composed of mHTT and other proteins, including transcription factors, ubiquitin, proteasomal subunits, and chaperones. It remains unclear whether the aggregates drive the disease, are merely coincidental, or serve a protective role by sequestering more toxic intermediates (reviewed in Jimenez-Sanches *et al*., 2017). Besides seeding protein aggregates, mHTT interacts with transcriptional regulators, causing epigenetic alterations, and changing gene expression (reviewed in Pogoda *et al*., 2021).

Multiple studies have shown that the HSR is impaired in HD models and involves reduced activity or levels of HSF1 (Chafekar *et al*., 2012; Riva *et al*., 2012; Maheswari *et al*., 2014; Das *et al*., 2016; Gomez-Pastor *et al*., 2017, reviewed in Gomez-Pastor *et al*., 2018). To restore HSR and HSF1 activity, small molecular drugs have been developed, including HSP90 inhibitors and HSF1 activators, giving various results in relieving HD symptoms (reviewed in Neef *et al*., 2011). HSP90 is an essential ATP-dependent chaperone, one of the cell’s most abundant proteins, and it interacts with a wide variety of clients (Taipale *et al*., 2012, Wei *et al*., 2024). The inhibitors disrupt HSP90’s ability to complete the chaperoning cycle, leading to increased tagging of clients to degradation (Solit and Chiosis, 2008). HSP90 inhibition also activates the HSR, which has been harnesses for therapeutic strategies (Whitesell *et al*., 2003; Waza *et al*. 2005; Neef *et al*., 2011). For instance, an HSP90 inhibitor HSP990 (Machajewski *et al*., 2007), was shown to induce HSF1 and temporarily relieve HD symptoms in mouse models. However, HSP990 did not give sustained results, likely due to altered chromatin landscape impairing the HSR (Labbadia *et al*., 2011). In agreement, synthetic glucocorticoid dexamethasone downregulated HSP90, induced HSF1, and transiently relieved HD symptoms in mouse and fly (Maheshwari *et al*., 2014). Also direct activation of HSF1 *via* plant derived Withaferin A ameliorated HD symptoms in mice (Joshi *et al*., 2021). However, induced HSR *via* HSF1 activation or by inhibiting HSP90 has been reported to provoke an earlier appearance of mHTT containing aggregates (Calamini *et al*., 2011; Bersuker *et al*., 2013). The various results, and the consistent lack of long-lasting relieve, manifest the difficulty to treat HD *via* small molecule inhibition. Moreover, the inability to ameliorate HD *via* induced HSR raises questions on how cells combat distinct proteotoxic stresses and which stress pathways could effectively mitigate polyQ stress.

To characterize cellular responses to distinct proteotoxic conditions, we compared transcription upon heat shock, HSP90 inhibition, and polyQ aggregation using existing information-rich RNA-seq datasets. First, we analysed polyA+ RNA expression in muscle (*quadriceps femoris*) of three mouse genotypes: wild type (*WT*), HD model *R6/2*, and *Hsf1* knock-out (*Hsf1^-/-^*), all subjected to heat shock or HSP90 inhibition (Neueder *et al*., 2017). The *R6/2* mouse expresses 115-150 Q repeats with a truncated (exon 1) mHTT, providing an excellent model for polyQ aggregation and fast HD progression (Mangiarini *et al*., 1996). Next, we assessed tissue-specific reprogramming of RNA synthesis using data from *Q175* HD mouse (Langfelder *et al*., 2016) that contains 175 Q repeat in full-length mHTT, and shows a late disease onset (Menalled *et al*., 2012). Finally, we addressed age-derived changes by investigating RNA expression in striatum of 2-month, 6-month, and 12-month-old *WT* and *Q175* mice (Diaz-Castro *et al*., 2019). We found heat stress, HSP90 inhibition, and polyQ aggregation to launch profoundly different RNA expression programs. Moreover, mice under chronic stress of either HSF1 deficiency or polyQ expression displayed a systemic inability to mount acute responses. Through decades, HSF1-driven chaperone expression has provided a robust model for induced transcription (Guertin *et al*., 2010) and fostered key insights in proteostasis (Hu *et al*., 2020). However, focusing on this one conserved pathway has shadowed the complexity of transcriptional responses. We found clear *Hsp* induction only upon acute stresses of heat shock and HSP90 inhibition, whereas mice carrying polyQ aggregates responded *via* RNAs involved in autophagy, repression of transcription, and chromatin remodeling. Simultaneously, key metabolic pathways and tissue-specific processes were repressed. We conclude HSR to constitute a variety of stress-specific responses, driven *via* distinct *trans*-activators and tailored to the proteotoxic condition. We propose that full understanding of the profoundly different RNA expression programs, involving tissues-specific pathways and stress-specific protection, is crucial for designing therapeutic strategies, and dissecting how cells combat adverse - acute or chronic - conditions. Finally, RNA expression programs across tissues highlight the importance of stage-dependent timing of HD treatments, and the lack of commonly induced or repressed genes suggest targeting pathways, such as energy metabolism, detoxifying enzymes, and autophagy, to ameliorate late-stage HD.

## Results

### Proteotoxic conditions trigger stress-specific RNA expression signatures

To characterize transcriptional responses to distinct proteotoxic stresses, we analysed RNA expression in muscle of *WT*, *R6/2*, and *Hsf1^-/-^* mice exposed to heat shock or HSP90 inhibition (Neueder *et al*., 2017). The heat shock *in vivo* was a whole-body heat pad for 20 minutes, followed by a 4-hour recovery (HS). For HSP90 inhibition, mice were treated with the small molecule HSP990 for 4h (i90). Respective control conditions were non-heat shocked (NHS) and vector control injected (iC) mice (Neueder *et al*., 2017). We downloaded the polyA+ RNA-seq libraries (Fig. S1A), mapped the raw data to mouse genome (mm10), and ensured high replicate correlation and quality (Fig. S1B). To find differentially expressed genes, we used DESeq2 that compares variance within replicates to the expression difference between conditions (Love *et al*., 2014). Differential expression revealed that heat shock, HSP90 inhibition, and polyQ aggregation launched remarkably distinct RNA expression programs (Fig 1A-B). Only five genes showed statistically significant induction in all three proteotoxic stresses, while no gene was significantly repressed in all stress conditions (Fig. 1B, Data S1). To understand the characteristics of transcription programs across genotypes and conditions, we performed principal component analysis (PCA), which projects high dimensional data into lower dimensions, maximizing the retained variance. Transforming the expression programs into principal components (PCs) 1 and 2 confirmed that distinct proteotoxic stresses directed RNA expression in remarkably different directions (Fig. 1C). While the polyQ stressed *R6/2* mice were separated from *WT* and *Hsf1^-/-^* mice along the PC1 (x-axis holding 33% of total variance), heat stress and HSP90-inhibition triggered cellular responses to opposite directions, visualized along the PC2 (y-axis, 12% of total variance). These results demonstrate the profoundly diverging responses depending on the proteotoxic condition.

**Figure 1.**
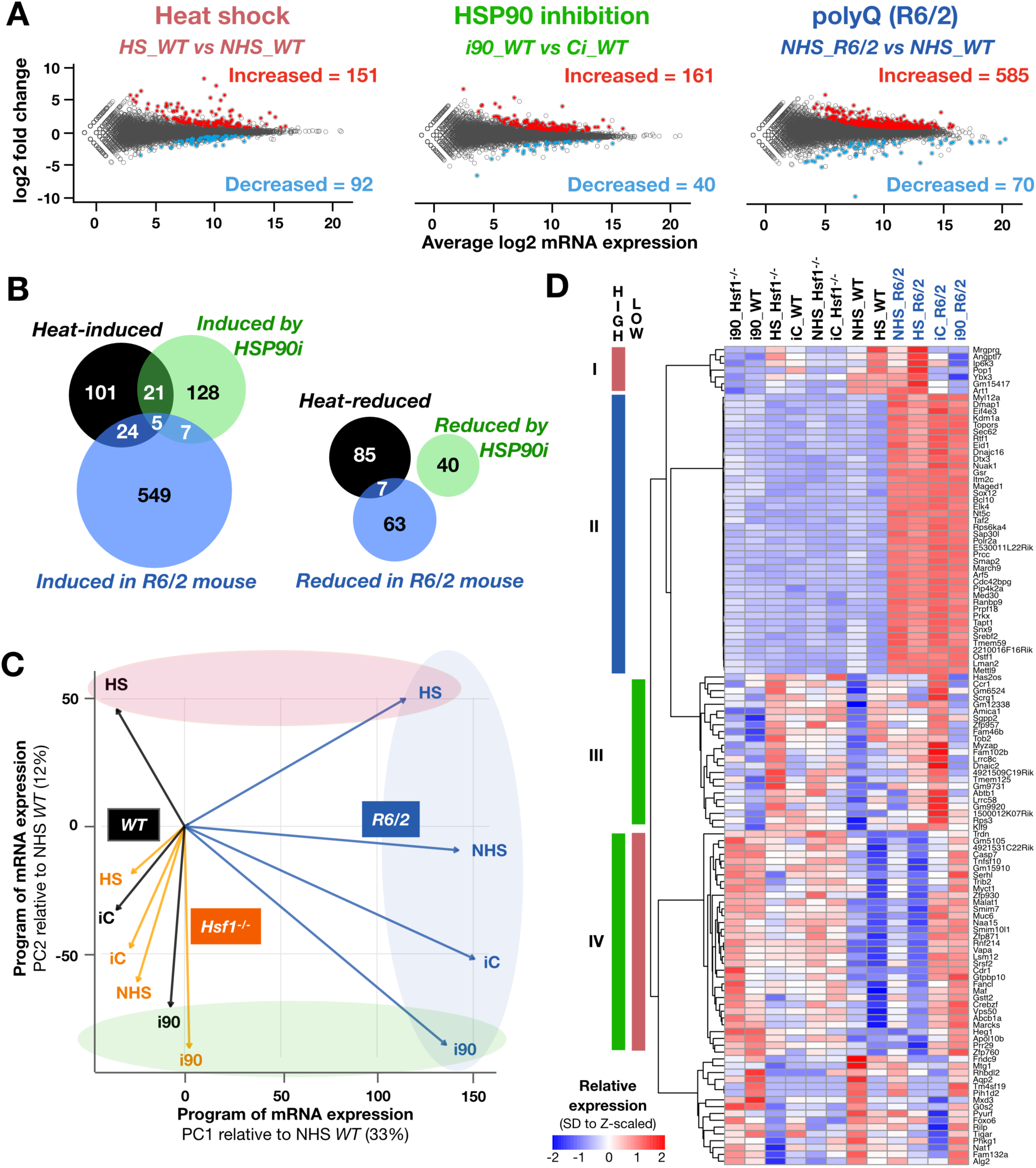
Heat shock, HSP90 inhibition or polyQ aggregation launch distinct RNA expression programs. **A)** MA-plots showing the average expression (x-axis) and stress-induced change (y-axis) of RNA expression. Left: heat shock (HS). Middle: HSP90-inhibition (i90). Right: polyQ expression *(R6/2).* Significantly increased (red) and decreased (light blue) RNAs were identified with DESeq2 (p-value < 0.001 and |fold change| 1.25. **B)** Venn diagram depicting the number of significantly induced (left) or repressed (right) genes. **C)** RNA expression program in *WT* (black), *R6/2* (blue) and *Hsf1^-/-^* (orange) mice under distinct proteotoxic stresses, projected to principal components 1 (x-axis) and 2 (y-axis). RNA expression program in non-stressed *WT* is positioned into the origo. The arrows show the direction and distance of transcription programs between phenotypes and conditions. PolyQ (*R6/2*), heat stress (HS), and HSP90 inhibition (i90) launch transcription programs into distinct directions, indicated with shaded blue, red, and green areas. **D)** RNAs with top variance compared in a heatmap. Clusters I-IV contain RNAs with high (left) or low (right) expression in HS, HSP90i, or *R6/2,* colored as the shading in panel C.

To understand which genes contributed most to the variance in stress-induced programs, we selected most important RNAs for PCs 1-3 and visualized their expression in a heatmap (Fig. 1D). As expected from the PCA (Fig. 1C), polyQ expressing mice showed a distinguishable RNA expression program, with several genes induced specifically in *R6/2* (Fig. 1D, group II). These genes were enriched with functions in negative regulation of RNA Polymerase II, histone phosphorylation coupled to DNA damage and chromatin remodeling, and lysosome activity (Data S2). A handful of genes (group I) were increased upon heat stress in *WT* and *R6/2*, but not in *Hsf1^-/-^*, mice (Fig. 1D). The only chaperone or co-chaperone among the RNAs carrying most variance was *Dnajc16*, which was highly expressed in *R6/2* mice (Fig. 1D) and regulates the size of autophagosome (Yamamoto *et al*., 2020).

### Acute stress induces chaperones – chronic polyQ stress activates autophagy

The profoundly different transcription programs in acutely (HS, HSP90i) and chronically (polyQ) stressed mice prompted us to seek genes that were induced (Fig. 2A-B) or repressed (Fig. S2B-C) across proteotoxic conditions. The five genes induced in all three stresses encoded inducible HSP90 (*Hsp90ab1*) and co-chaperone DNAJA4, which has been described both as a heart-enriched co-chaperone for HSP70 (Abdul *et al*., 2002; Hafizur *et al*., 2006) and a membrane-associated protein implied in cholesterol biosynthesis (Robichon *et al*., 2006). The five commonly induced genes also encoded solute carrier SLC7A5 that transports amino acids across membranes (Kanai *et al*., 1998), and Aldehyde oxidase 1 (AOX1), which neutralizes aldehydes and heterocyclic compounds (Wright *et al*., 1997).

**Figure 2.**
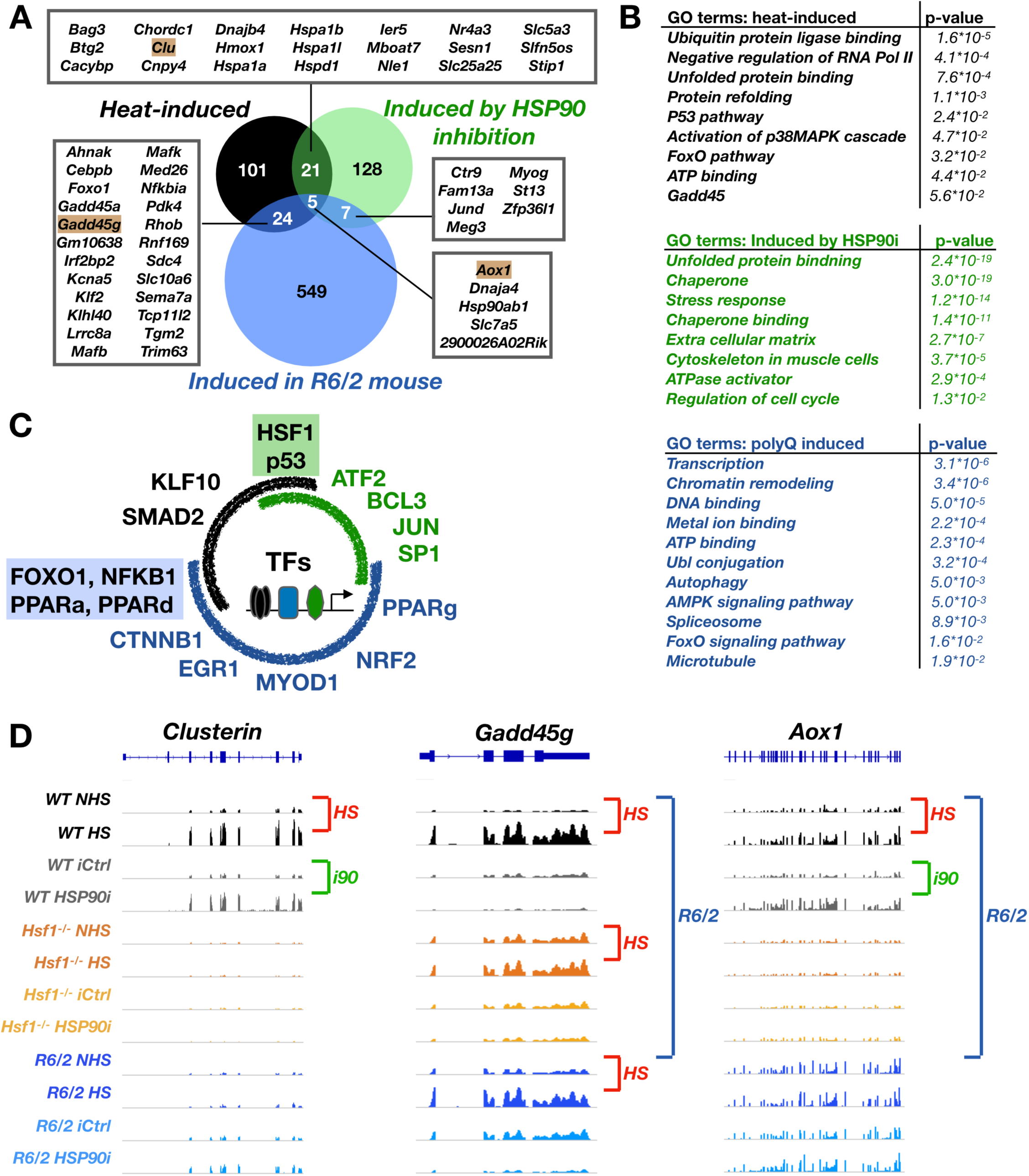
Heat stress and HSP90 inhibition induce chaperones, polyQ aggregation increases lysosome and autophagy related RNAs. **A)** Significantly increased RNAs compared in a Venn diagram. Genes that are induced in two or three proteotoxic stress conditions are listed in boxes. Genes indicated with golden background are shown in browser graphs in panel D. **B)** Enriched GO terms among heat-induced, HSP90i-induced, and polyQ-induced genes. FDR corrected p-value (Benjamini) is indicated for each category. **C)** Enrichr analysis of transcription factors driving stress-specific transcription programs. Black, green, and blue arches indicate TFs associated with heat, HSP90i, and polyQ induced RNAs, respectively. Overlapping arches and shaded text show factors associated two stress conditions. Complete enrichr report is in Data S3. **D)** Genome-browser examples of genes induced in a proteotoxic stress-specific manner. Clusterin *(Clu)* and *Gadd45g* mRNAs are induced in two conditions, *Aox1* is induced in all three conditions.

Beyond the five commonly induced genes, 21 were activated upon heat shock and HSP90 inhibition (Fig. 2A). These genes encode canonical HSP70 (*Hspa1a*, *Hspa1b*, *Hspa1l1*) and HSP60 (*Hspd1*) foldases, extracellular chaperone (*Clu*), co-chaperones (*Bag3*, *Chordc1*, *Dnajb4*, *Stip1*), and other known stress-induced proteins (*Cacybp*, *Ier5*, *Hmox1*). Moreover, genes encoding solute carriers across plasma (*Slc5a3*) and mitochondrial (*Slc25a25*) membranes were induced upon heat shock and HSP90 inhibition. While chaperone induction was abundant upon the acute stresses of heat shock and HSP90 inhibition, only three genes for HSP chaperone complexes (*Hsp90ab1*, *St13*, and *Dnajc18*, an ER localizing paralog of *Dnajc16*) were significantly activated in *R6/2* mouse (Data S1). The apparent lack of *Hsp* and *DnaJ* induction in muscle tissue of polyQ mice indicates that HD mice are either incapable of inducing chaperone machineries, or use alternative pathways for maintaining proteostasis, likely due to polyQ aggregates being too large for chaperone-mediated clearance (Cuervo and Wong, 2014). Indeed, *R6/2* mice expressed significantly elevated levels of RNAs involved in autophagy (15 RNAs) and lysosome (21 RNAs) pathways (Fig. 2B and Data S2), supporting polyQ aggregates to activate membrane-associated degradation, required for clearing large protein assemblies and damaged organelles.

### PolyQ containing mice remodel energy metabolism and reduce muscle-specific RNAs

Inhibition of HSP90 is expected to disrupt proteostasis primarily *via* its cytosolic client proteins. In contrast, heat stress holds broad potential to instantly break intermolecular bonds, and polyQ aggregates cause a cascade of disrupted cellular functions. Intriguingly, HSP90 inhibition reduced expression of its largest client group, kinases (Taipale *et al*., 2012), and induced RNAs for protein-centric functions, including chaperones and cytoskeletal proteins (Figs. 2B and S2B, Data S2). Heat stress and polyQ aggregation, instead, increased RNA expression for regulators of transcription, genome integrity, and metabolism, such as Growth arrest and DNA damage inducible 45 (GADD45), Forkhead box O1 (FOXO1), and their known genomic targets (Fig. 2A-B). Noteworthy is that the main repression in polyQ expressing muscle involved muscle-specific pathways, including myofibril, muscle contraction, and sarcoplasmic reticulum (Fig. S2B). The *R6/2* mouse also showed reduced levels of RNAs for essential metabolic processes, suggesting severely affected cell type-specific functions and cellular metabolism.

### Distinct sets of transcription factors drive proteotoxic stress responses

The different responses prompted us to analyse which transcription factors control the stress-specific changes in RNA expression. We performed Enrichr analyses (Xie *et al*., 2021) on transcription regulatory relationships (Han *et al*., 2018) and, as expected, found HSF1 as the most prominent *trans*-activator for genes induced upon heat shock and HSP90 inhibition (Fig. 2C, Data S3). Additionally, Tumor Protein 53 (p53), the coordinator of genome maintenance, cell cycle, and cancer suppression, was activated upon heat shock and HSP90 inhibition. No overlap on transcription factor activation was found upon HSP90 inhibition and polyQ expression. Instead, RNAs induced both in *R6/2* mice and upon heat shock were enriched for genomic targets of FOXO1, Nuclear factor kappa B precursor (NFKB1), and Peroxisome proliferator activated receptors (PPARs) (Fig. 2C, Data S3). Nf-KB is a key regulator of inflammatory responses (Pereira and Oakly, 2008), while FOXO1 is activated by nutrient deprivation and regulates metabolism and autophagy (Gross *et al*., 2009; Oli *et al*., 2021). Furthermore, targets for all the main PPAR-family members (PPAR-a, PPAR-d, PPAR-g), which coordinate fatty-acid oxidation and energy metabolism (reviewed in Zhao *et al*., 2024), were induced in *R6/2* mice (Fig. 2C).

For stress-repressed genes, no shared transcription factor was identified (Fig S2C). In polyQ expressing muscle, the downregulated genes contained targets of Krüppel like factor 3 (KLF3), which functions primarily as a transcription repressor, and controls key muscle genes (Himeda *et al*., 2010; Pearsson *et al*., 2011). Additionally, genomic targets of Sterol regulatory element binding factor 1 (SREBF1) and Carbohydrate responsive element binding protein (ChREBP) were repressed (Fig. S2C). While ChREPB promotes conversion of excess carbohydrates to triglycerides (Iizuka and Horikawa, 2008), SREBF1 controls cholesterol and fatty acid synthesis (Shimano, 2001; Horton *et al*., 2002). These changes in RNA expression lend strong evidence that *R6/2* mice reduce glycolysis (repressed ChREPB targets), experience nutrient deprivation (FOXO1 activation), suppress fatty-acid synthesis (repression of SREBP1), and activate fatty-acid oxidation and lipid catabolism (activated PPAR targets) as an alternative energy source. Altogether, the adapted RNA expression in *R6/2* mouse points to a profound shift energy in metabolism, from lipid biosynthesis to lipid catabolism.

### 3D Stress Cube: Gene-specific analyses of global responses

DEseq2 uses strict statistical criteria to call differentially expressed genes, which could miss key genes and patterns in RNA expression programs. To provide gene-centric views within the context of the global changes, we generated an interactive 3D visualization tool, termed 3D Stress Cube (Fig. 3 and File S1). This easy-to-use tool positions each RNA along x, y, and z axes according to log2 fold changes (log2FC) upon heat shock, HSP90 inhibition, and polyQ expression (Fig. 3A). Each RNA is colored based on the directions of stress-induced changes (octants, Fig. 3B), allowing instant identification of gene-by-gene changes within transcriptional responses. As examples, *Foxf1* (octant 1, red) and *Ucp1* (octant 7, purple) were induced and repressed, respectively, in all three stress conditions (Fig 3A). RNAs with stress-specific responses are exemplified with *Vtcn1*, *Rab9b*, *Nccp*, *Angptl8*, *Odf3l2*, and *Ddit4* (Figs. 3A and S3). These analyses showed 17% of RNAs to be increased in all, and 8% to be repressed in all stresses (Data S4, all expressed RNAs, n=11,757), which indicate slightly higher and lower occurrences, than expected with a random change (12.5%). For RNAs with a total log2FC > 1.5 (Fig. 3A, n=3,496), the fraction of RNA induced in all stresses increased to 25%, and while the fraction of repressed in all remain below 8%. The cube is ready-to-use after downloading the File S1 in github.com/Vihervaara/3D-Stress-Cube and opening the html file in a web browser. Rotation, zooming, and information on each RNA (Fig. 3C) are provided. For manageable browsing, the cube contains genes that show a minimum of 1.5 total log2FC across the three stresses (n=3,496). Data S4 lists all expressed genes and their responses.

**Figure 3.**
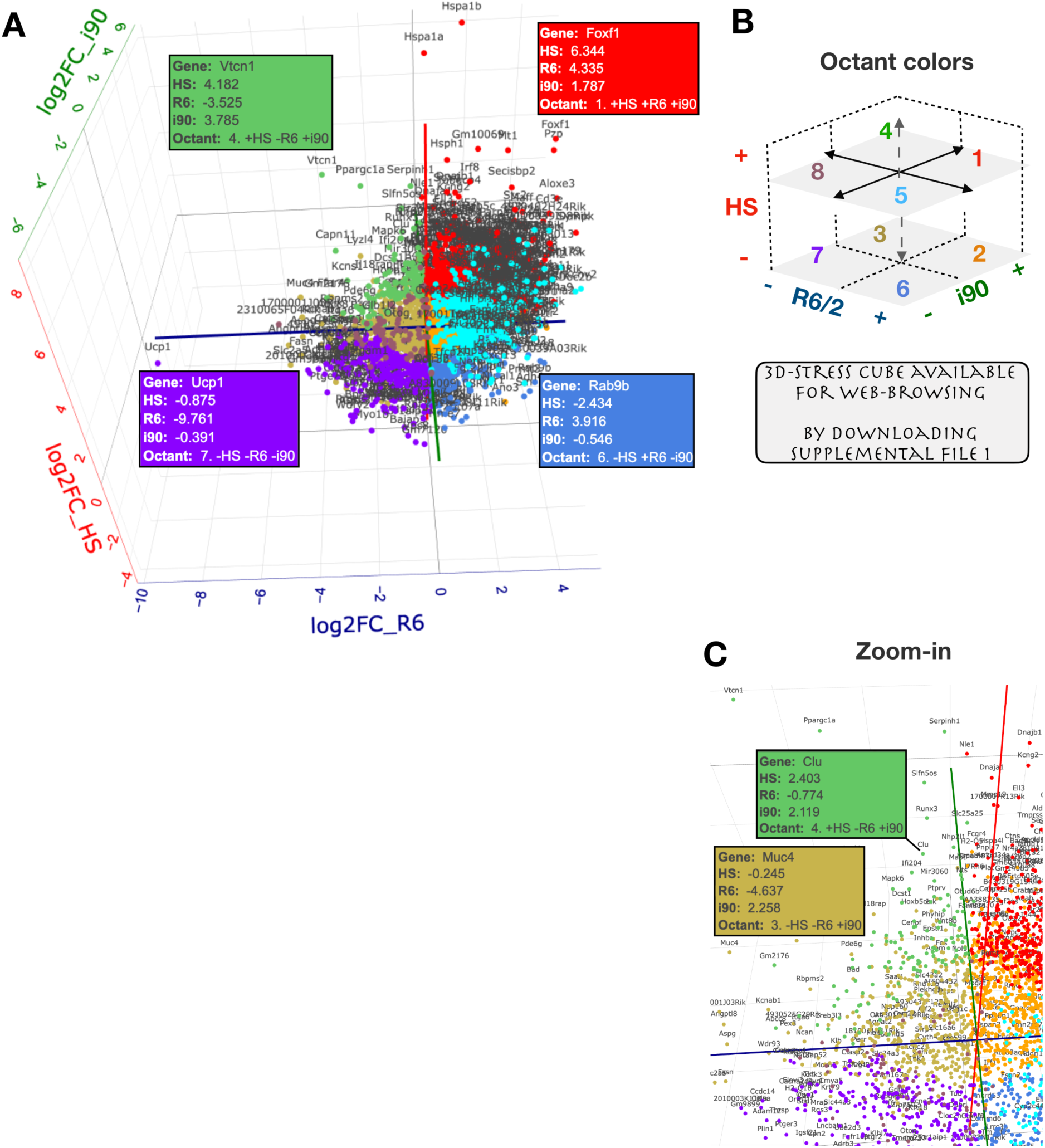
3D Stress Cube: Interactive visualisation of stress-induced transcriptional changes. **A)** Interactive 3D viewer shows transcriptional changes upon heat shock (HS), HSP90 inhibition (i90), and polyQ aggregation (R6). The RNAs are coloured based on the octant, i.e. the direction of the stress-induced changes, schematically shown in panel B. *Vtcnl* (green), *Ucp1* (purple), *Foxfl* (red) and *Rab9b* (matt blue) are highlighted as RNAs in the distinct octants, and the direction (– or +) and log2FC in each stress condition shown in the infobox. **B)** Color key for the octants. See Supplemental Figure 3 for example RNAs in octants 2, 3, 5 and 8. The 3D Stress cube is available for browsing by downloading and unzipping the File S1 and opening the .html file in a web browser. **C)** The cube allows zooming into specific areas, the boxes with RNA- specific information appear by placing the cursor over the dot. The 3D stress-cube was generated with plotly (Sievert, 2020) and stress-induced changes were counted as follows. Heat shock: log2(HS_WT/ NHS_WT). HSP90i: log2(i90_WT/iC_WT). PolyQ: log2(NHS_R6/ NHS_WT). The interactive cube contains polyA+ RNAs with combined log2FC >1.5 (n=3,496), counted as sqrt(log2FC_HS^2 + log2FC_R6^2 + log2FC_i90^2) >1.5. Data S4 contains all transcripts and their stress-induced expression changes.

### Systemically hampered responses in mice under chronic stress

To understand how mice under chronic stress mount transcriptional responses, we measured polyA+ RNA expression in *R6/2* and *Hsf1^-/-^* mice subjected to heat shock (Fig. 4) or HSP90 inhibition (Fig. 5). While *R6/2* mice are under chronic stress due to polyQ aggregation, *Hsf1^-/-^* mice have impaired metabolic pathways (Ma *et al*., 2015; Yan *et al*., 2002), indicative of chronic stress. Moreover, *Hsf1^-/-^* cells and mice are highly sensitive to induced stress (McMillan *et al*., 1998; Zhang *et al*., 2002). DESeq2 analyses showed a drastic reduction in differentially expressed genes upon heat shock in *R6/2* and *Hsf1^-/-^* mice (Figs. 4A, 5A and S4), as compared to *WT* mice (Fig. 1A). These results are well inline with the original analysis by Neueder *et al*. (2017), where blunted responses in polyQ mice were reported. Importantly, the limited stress responses included global induction and repression, and extended well beyond HSF1 target genes (Figs. 4A-B, 5A-B).

**Figure 4.**
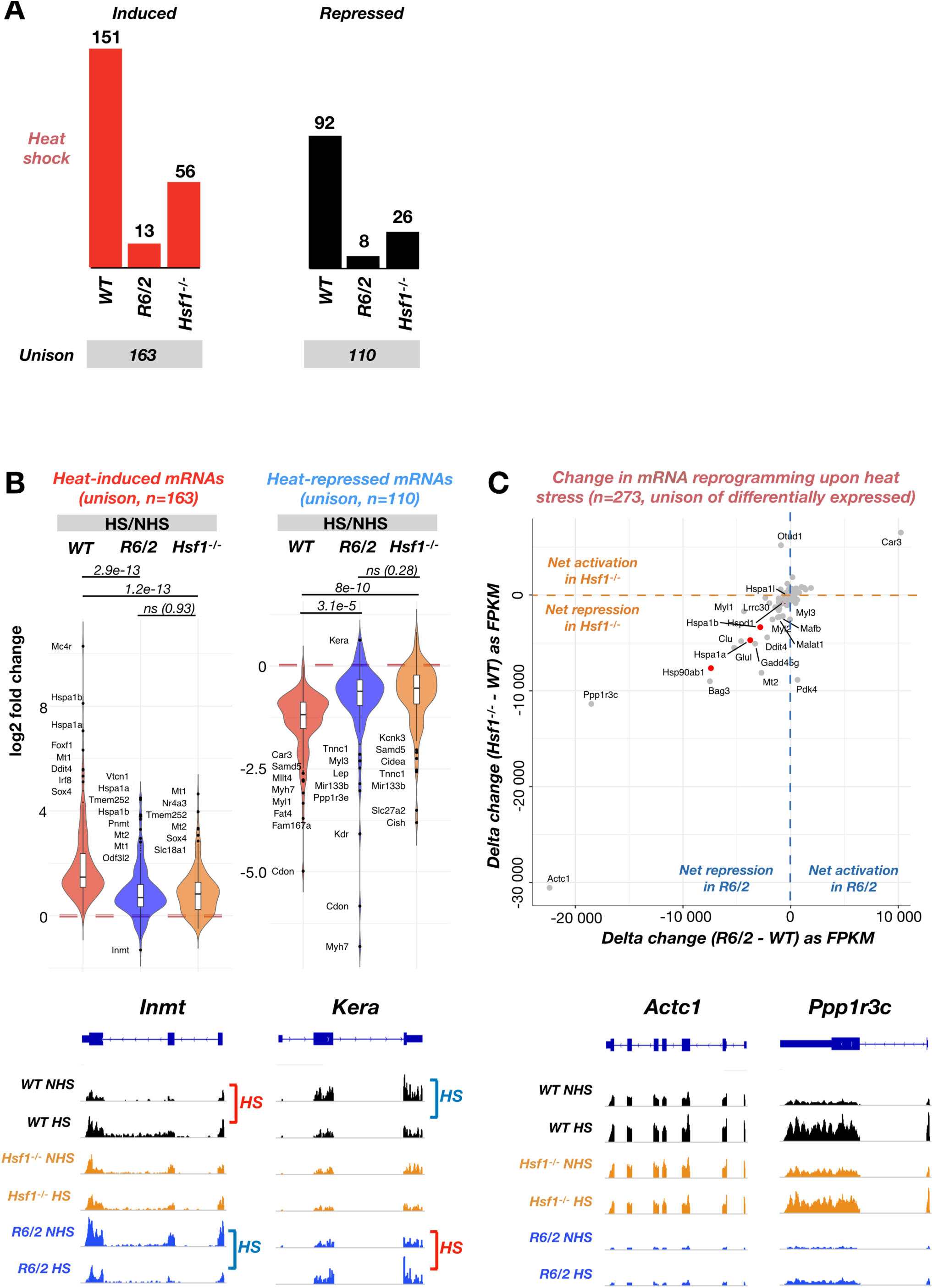
Heat shock response is systemically blunted in mice under chronic stress. **A)** Number of differentially expressed RNAs upon heat shock in *WT*, *R6/2,* and *Hsf1^-/-^* mice. Unison of induced (left) and repressed (right) RNAs are indicated below the bar charts. Related Supplemental Figure 4A contains MA-plots for *R6/2* and *Hsf1^-/-^* mice. **B)** Upper: Log2 fold change of differentially expressed RNAs upon heat shock, analysed for the unison of induced (left) and repressed (right) RNAs in the three mouse genotypes. P-values from paired student’s t-tests are indicated for each pairwise comparison. Lower: Genome browser visualisation of RNA expression from two outlier genes, *Inmt* and *Kara,* showing rare examples of changed directionality of stress-response in *R6/2* mouse. **C)** Upper: Acute stress response compared in WT *versus* chronically stressed *(R6/2* or *Hsf1^-/-^)* mice. Delta change in FPKM is derived as HS - NHS, and compared as *R6/2 -* WT (x-axis) or *Hsf1^-/-^ WT* (y-axis). Lower: Genome browser examples of two outlier genes, *Actc1* and *Ppp1f3c*.

**Figure 5.**
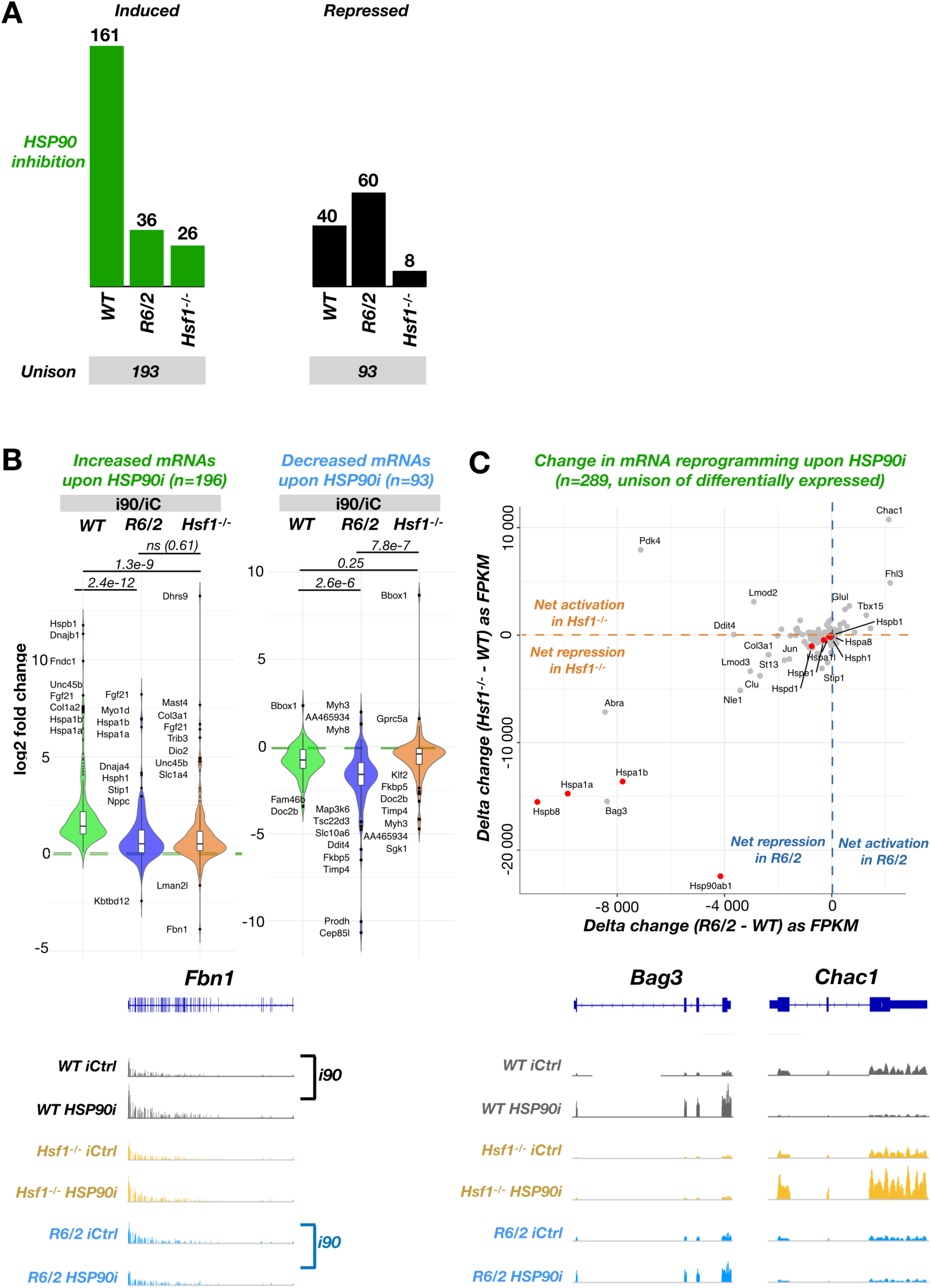
Transcriptional response to HSP90 inhibition is systemically blunted in mice under chronic stress. **A)** Number of differentially expressed RNAs upon HSP90 inhibition in *WT, R6/2,* and *Hsf1^-/-^* mice. Unison of induced (left) and repressed (right) RNAs are indicated below the bar charts. See related Supplemental Figure 4B for MA-plots of *R6/2* and *Hsf1^-/-^* mice. **B)** *Upper:* Log2 fold change of differentially expressed RNAs upon HSP90 inhibition, analysed from the unison of induced (left) and repressed (right) RNAs in the mouse genotypes. P- values from paired student’s t-tests are indicated for each pairwise comparison. *Lower:* Genome browser example of an outlier gene, *Fbn1.* **C)** *Upper:* Acute stress response in *WT versus* chronically stressed (*R6/2* or *Hsf1^-/-^)* mice. Delta change in FPKM is derived as HSP90i - Ci, and compared as *R6/2 - WT* (x-axis) or *Hsf1^-/-^ WT* (y-axis). *Lower:* Genome browser examples of two outlier genes, *Bag3* (left) and *Chac1* (right).

Since fold changes can be large for genes with a low RNA expression, we compared gene-by-gene expression as difference (stress - control) between *WT* and chronically stressed mice (Figs. 4C and 5C). This gene-centric comparison revealed a striking similarity between *R6/2* and *Hsf1^-/-^* mice, as genes with reduced induction in polyQ mice were also less induced in mice that lacked HSF1 (Figs. 4C and 5C). Given the different transcription programs in *Hsf1^-/-^ versus R6/2* mice (Fig. 1B-C), these results indicate a systemic inability to mount acute transcriptional responses in chronically stressed mice, whether due to polyQ aggregates or lack of HSF1. This inability comprehended induction and repression, occurred upon heat shock (Fig. 4) and HSP90 inhibition (Fig. 5), and included HSF1-dependently and independently activated genes. Consequently, severely and broadly dampened ability to mount acute stress responses emerged as a feature of chronically stressed mice, not a limited inactivity of a single transcription activator or a pathway.

### Transcriptional changes to polyQ expression are tissue-specific

The distinct transcription programs (Figs. 1-2) and stress responses (Figs. 3-5) in striatal muscle of *WT* and *R6/2* mice prompted us to examine RNA expression across tissues. For this, we downloaded raw RNA-seq data generated by the HDinHD consortium (Langfelder et al., 2016; hdinhd.org) in 11 tissues of *WT* and *Q175* mice. Particularly, we examined whether polyQ aggregates caused similar transcriptional changes across tissues, and whether the absence of the canonical HSP response to polyQ aggregation was a general feature of HD pathology. PCA revealed profoundly tissue-specific RNA expression in *WT* and *Q175* mice (Fig. 6A). While brain tissues were clustered close to one another (PCs 1 and 2), skeletal and heart muscle were separated from adipose tissues and skin along the PC2 (Fig. 6A). Importantly, differences in RNA expression programs between *WT* and *Q175* mice were minute as compared to the tissue-specific expressions (Fig. 6A). These results indicate that the adaptation to polyQ aggregates occurred within tightly tissue-constrained transcriptional environments. Next, we zoomed into the brain regions and found each region to exhibit a separable transcription profile (Fig. S5A). The RNA expression variance attributable to polyQ aggregates, however, remained very small (Fig. S5A). Thus, transcriptional responses to polyQ aggregates were remarkably tissue-specific, even within the central nervous system. Accordingly, DESeq2 analysis found hundreds of differentially expressed genes in the brain regions of *Q175* mouse, as compared to *WT* mouse, but little overlap in polyQ-induced changes between tissues (Figs. 6B, S5B). GSEA analysis of affected pathways revealed repressed genes in *Q175* mice to be enriched with cell type-specific functions and energy metabolism (Data S5). These results are well in-line with the reduced expression of RNAs for muscle-specific functions and metabolic pathways in *R6/2* mouse muscle (Figs. 2 and S3). Conversely, immune related RNAs were upregulated in several brain regions (most notably brainstem and corpus callosum) and in adipose tissues, while they were suppressed in heart and gastrocnemius muscle (Data S5). We did not find evidence for the canonical HSP-inducing stress response in any of the investigated tissues of *Q175* mouse (Data S5).

**Figure 6.**
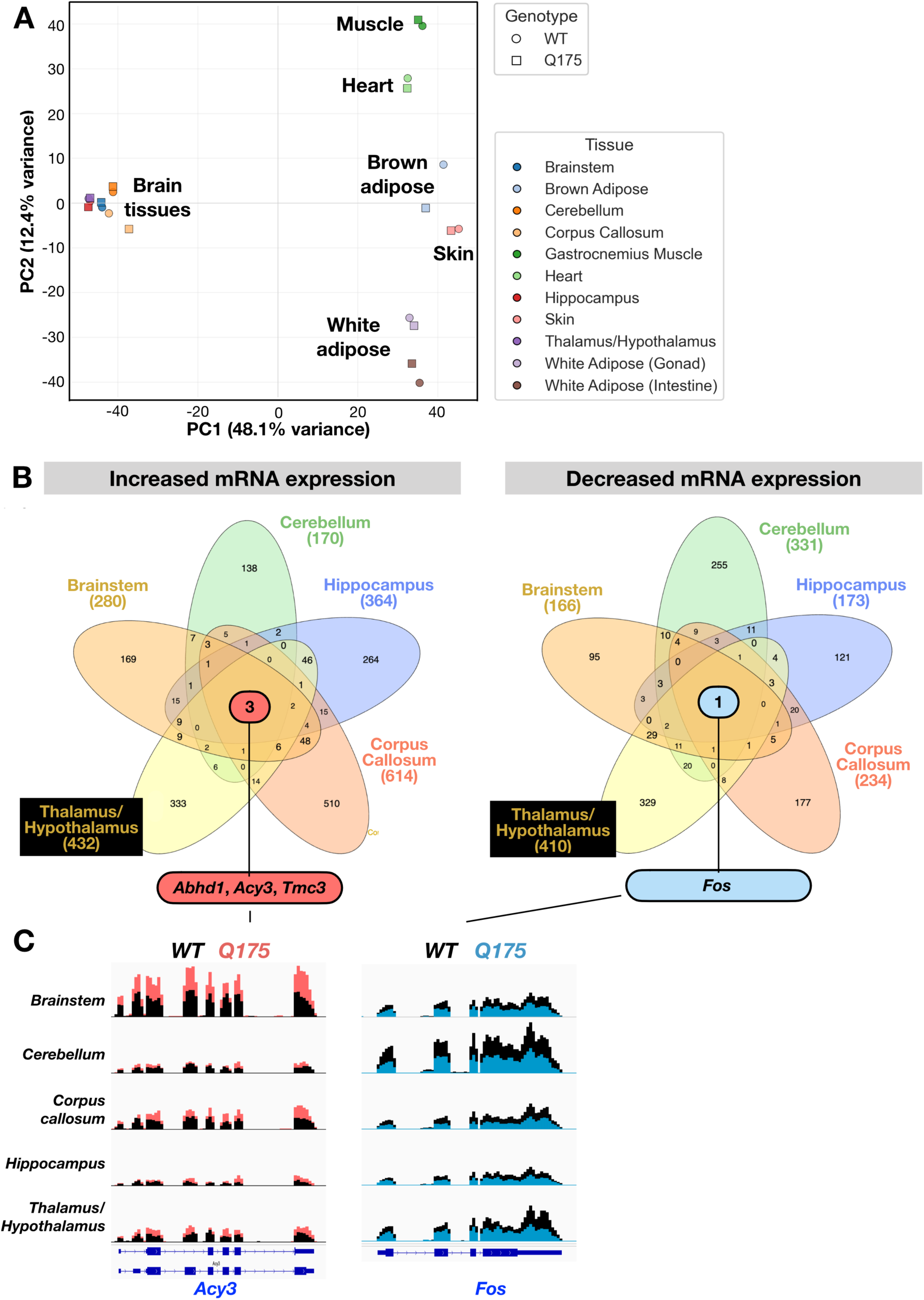
PolyQ stress causes tissue-specific reprogramming of RNA expression. **A)** PCA analysis of RNA expression across tissues of *WT* and *Q175* mice. **B)** Comparison of differentially increased (left) and reduced (right) RNAs between *WT* and *Q175* mice across tissues. Differential expression was analysed with DESeq2 using p-value < 0.05 and |log2FC| > 0.5 as thresholds. The thee RNAs that are differentially induced *(Abhd1, Acy3, Tmc)* and the one *(Fos)* that is differentially reduced across brain tissues are shown. **C)** *Acy3* (left) and *Fos* (right) RNA expression across brain tissues.

Across the brain regions analysed, only three mRNAs (*Abdh1*, *Acy3*, *Tmc3*) were consistently upregulated, and one (*Fos*), consistently downregulated in *Q175* mouse (Fig. 6B-C). *Fos proto-oncogene* is a rapidly responding gene that encodes c-Fos, a component of a potent *trans*-activator, Activator protein 1 (AP1). In neurons, reduced c-Fos levels are associated with impaired synaptic input, and altered c-Fos expression has been reported in some HD models (Murphy *et al*., 1991; Rawat *et al*., 2016). Among the consistently induced RNAs, *Aminoacylase 3* (*Acy3*) is an enzyme involved in detoxification, capable to hydrolyze acylated amino acids generated as metabolites of protein degradation (Pushkin *et al*., 2004). α/β-hydrolase fold containing 1 (*Abdh1*) is likely an enzyme with lysophilic lipase activity and involved in stress responses (Edgar 2003; Lord *et al*., 2013). Transmembrane channel like 3 (*Tmc 3*) encodes a poorly characterized ion-channel, expected to function in mechanosensing (Braz *et al*., 2025). Notably, *Abdh1* was previously reported to be upregulated in an HD cell model (van Roon-Mom *et al*., 2008), which together with its increased expression in brain regions of *Q175* mouse could reflect tissue-wide compensatory mechanisms to maintain lipid homeostasis. Although *Abhd1* and *Tmc3* emerge as potential markers and influencers of HD pathology, both genes partially overlap with a strongly, and seemingly ubiquitously, expressed gene (*Preb1* and *Gm16638*, respectively) in the mouse genome (Fig. S5C). These overlaps call for strand-specific quantification of RNA expression to ensure correct quantification of RNA levels. To this end, *Acy3* and *Fos* provide robust and unambiguous examples of consistently increased and repressed RNAs across HD brain tissues (Fig. 6C), suggesting their potentiality as HD markers and therapeutic targets. We conclude that similarities in responses to polyQ aggregates were more evident in induced and repressed pathways, rather than in differentially expressed genes. These findings underscore that transcriptional adaptation to polyQ toxicity is deeply tissue-specific, and calls diagnostic and treatment strategies to target processes, such as energy metabolisms and lipid homeostasis, rather than individual proteins.

### Ageing changes RNA expression in mouse striatum but has a limited effect on *Hsp* levels

The cells’ ability to respond to proteotoxicity changes with age. To analyse whether the lack of increased *Hsp* mRNA expression in polyQ mice was age-dependent, we re-mapped data by Diaz-Castro *et al*. (2019), generated in striatum of *WT* and *Q175* mice of 2, 6, and 12 months of age. As reported in the original study (Diaz-Castro *et al*., 2019), we identified several genes whose expression was significantly induced or repressed in *Q175* mice during ageing. Intriguingly, these genes included the induction of *Acy3* (Fig. 7A), *Tmc3* and *Abdh1* (Fig. S6A), and repression of *Fos* (Fig. 7B), further strengthening their potential as tissue-wide markers of HD. Comprehensive analysis of *Hsp* and *Dnaj* RNAs, instead, identified only modest differences in *WT* versus *Q175* striatum. These changes included reduced expression of *Hsph1,* and increased expressions of *Hspa8* (HSC70) and *Dnajc18* mRNAs (Fig. 7C-D). We found no clear pattern of induction or repression for components shared by chaperone complexes (Figs. 7C-D, S6B and S7A), or *Hsf1* (Fig. S7B) in ageing striatum. Based on RNA expression programs across tissues, we conclude that polyQ stress reduces expression of tissue-specific RNAs, alters energy metabolism, and has limited or varying effects on genes involved in canonical acute stress response. The analyses of RNA expression across tissues highlight energy metabolism pathways as potential targets and causes for impaired cellular functions in polyQ diseases. Moreover, the results suggest activities of key transcription factors, including FOXO1, GADD45, PPAR and cFos, and enzymes, such as ABHD1 and ACY3 as markers and contributors in HD pathology.

**Figure 7.**
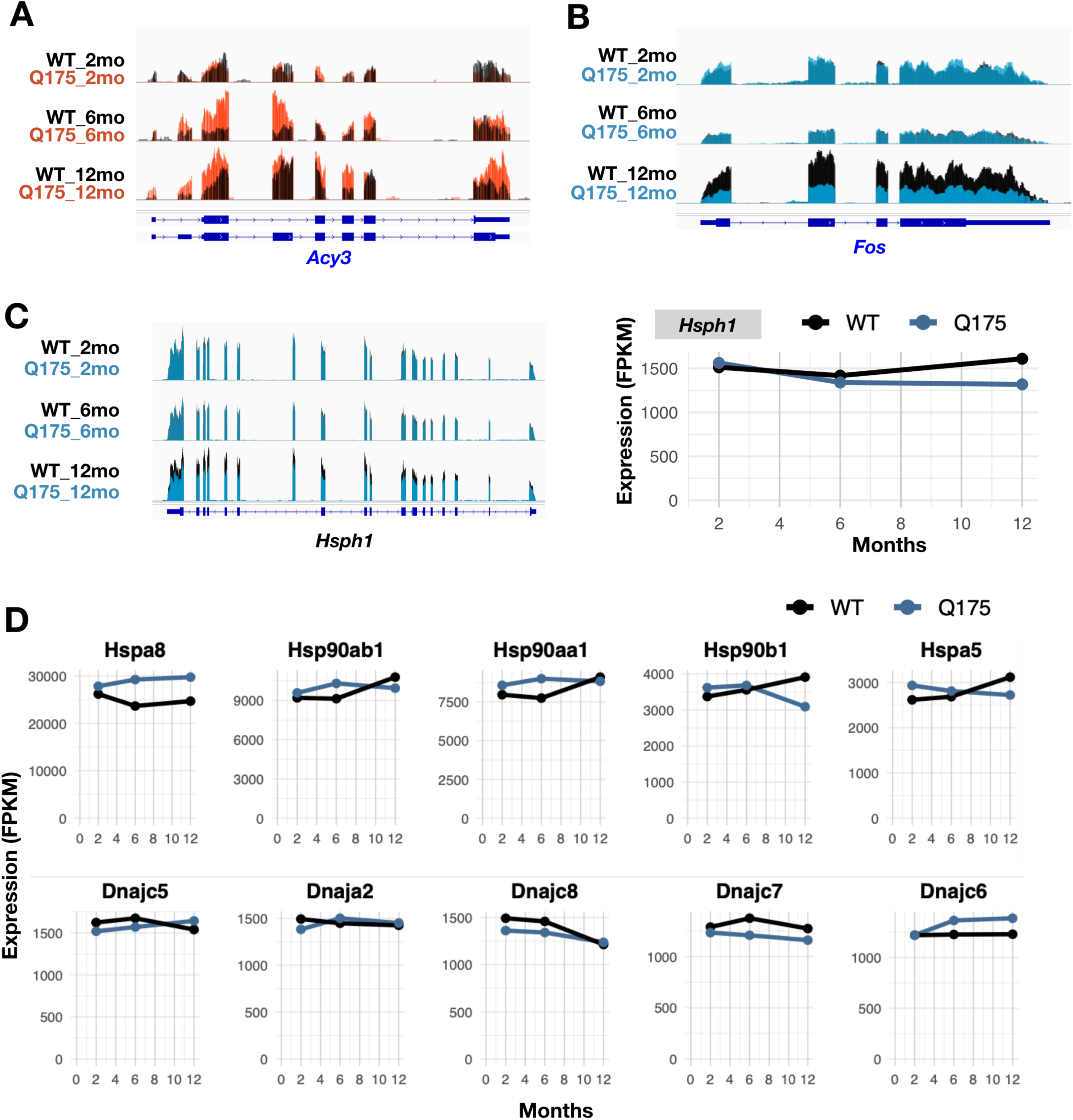
RNA expression changes in striatum of ageing *WT* and *Q175* mice. **A-C)** Genome browser views on mRNAs whose expression is **A)** increased *(Acy3)* and **B)** decreased *(Fos)* in ageing *Q175* mouse. **C)** *Hsph1* mRNA expression shows a modest decline in ageing striatum of *Q175* mice. **D)** Expression changes of the highest expressed *Hsp* (upper panels) and *Dnaj* (lower panel) mRNAs in striatum of *WT* and *Q175* mice. For expression changes of all other *Hsp* mRNAs and most (FPKM > 200) *Dnaj* mRNA, see related Supplemental Figures S6B and S7A, respectively.

## Discussion

### The HSR comprises several stress-specific transcription programs

Since the discovery of heat shock proteins (Lindquist 1980) and HSF1 as their heat-induced *trans*-activator (Wu *et al*., 1987), analyses of stress responses have focused on this central and conserved pathway. As a result, many stress conditions that increase HSP expression, including heavy metals, infections, and neurodegenerative diseases, have been coined to induce the HSR (Morimoto, 2008). With the advent of techniques that identified all HSF1 binding sites, or tracked transcription and RNA expression genome-wide, a remarkable complexity of HSF1 in controlling promoters and enhancers, and driving distinct transcription programs in stress, cancer, and differentiation, emerged (Hahn *et al.,* 2004; Åkerfelt *et al.,* 2010; Guertin and Lis, 2010; Mendillo *et al.,* 2012; Santagata *et al*., 2013; Vihervaara *et al*., 2013; 2017; 2021; Scherz-Shouval, 2014; Duarte *et al*., 2016; Mahat *et al*., 2016; Himanen *et al*., 2022). In this study, we took a genome-wide approach to compare RNA expression changes triggered by distinct protein damaging conditions. We found proteotoxic stresses, which all could be categorized under the broad term of HSR (Morimoto, 2008), to launch RNA expression programs to remarkably distinct directions (Fig. 1). Only few genes were statistically significantly induced or repressed in two or three of the stress conditions (Fig. 2). Even with loose criteria - analysing directions of the responses - less than a third of genes were either induced (25%) or repressed (8%) in all three proteotoxic conditions (Fig. 3A and Data S4). We highlight that HSR should be considered as an umbrella term for a multitude of transcription programs that drastically differ from one another - each tailored to combat the specific adverse condition.

### Proteotoxic-specific responses occur *via* distinct transcription factors

Cellular stresses, such as oxidative stress, hypoxia, ER load, and temperature changes, activate dedicated stress-specific *trans*-activators (reviewed in Vihervaara *et al*., 2018). However, no genome-wide comparison of transcription factor activities under distinct proteotoxic stresses had been performed. We found HSF1 and p53 targets to be induced upon acute stresses. Instead, the chronic stress of polyQ aggregation activated FOXO, PPAR, Catenin beta (CCNTB1) and Myoblast determination (MYOD1) targets, and repressed lipid and glucose metabolism genes of SREBF1 and ChREPB (Fig. 2). Altogether, the proteotoxic stress responses were driven *via* separate set of transcription factors. The main overlap, i.e. the HSF1-activated chaperone network, covered less than 10% of induced genes upon heat shock or HSP90 inhibition (Fig. 2). While acute stress induced chaperones, polyQ aggregates increased RNAs for autophagy and transcription repression, indicating a shift from RNA production to protein degradation. Moreover, polyQ expression repressed RNAs involved in tissue-specific functions and energy metabolism, further pointing to a broad slow-down in basal cellular processes in HD mice. Particularly, changing from glucose usage and storage to lipid catabolism (Figs. 2 and S2), and disrupted mitochondrial processes (Data S5), indicate HD mice to struggle to meet the energy requirements. Worth noting is that chaperoning *via* foldases, such as HSP70, HSP90, TRiC, HSP10-HSP60, and HSP110, is highly ATP-consuming. Likewise, maintenance of the acidic environment in lysosomes, and large-scale membrane remodeling in autophagy, requires considerable amount of energy. Taken together, the RNA expression changes across tissues of HD mouse manifest reduced basal operations, and attempt to maintain proteostasis *via* autophagy, likely in an environment with a limited energy supply.

### Impaired acute stress responses emerge as a hallmark of constitutive stress

Mice under chronic stress, either due to polyQ aggregates or lack of HSF1, displayed severely limited responses to acute stress (Figs 4 and 5). Intriguingly, RNA expression programs in HD mice suggest reduced RNA Polymerase II activity, a condition that ablates transcriptional responses to acute stress and differentiation (Prajapati *et al*., 2024). Many attempts have been made for ameliorating HD symptoms through restoring the HSR, which relies on the ability to launch a transcriptional response. Analyses of RNA expression programs in *R6/2* and *Q175* HD mice demonstrated profound challenges in inducing HSR in late-stage HD; First, the systemic deficiency to mount acute stress responses severely limits induced chaperone synthesis (Figs. 4-5). Second, repression of RNA Polymerase II in HD mice (Figs. 1-2 and S2, Data S5), could constitute the underlying cause for ablated responses, and impair any *trans*-activator driven change. Third, no or low HSP increase was detected across polyQ expressing tissue in *Q175* mice of distinct ages (Figs. 5-7, Data S5), pointing to cells preferring other means for restoring proteostasis in HD stages with solid-stage aggregates. These alternatively pathways could involve autophagy and detoxifying enzymes, such as ACY3 and AOX1 (Figs. 2, 6-7). Fourth, the environment with altered metabolism and presumably reduced energy availability challenges activation of energy-heavy processes, including transcription, protein re-folding, and aggregate clearance. Indeed, support for energy production could aid HD cells to induce transcriptional responses and sustain cell-specific functions. We propose that systemic inability to mount acute stress responses is a hallmark of chronic stress, and involves globally reduced transcription and disturbed energy balance.

### Emerging strategies for *mHtt* silencing, pathway targeting, and treatment timing

Recent years have witnessed an expansion of therapeutic strategies for HD treatment, including genome editing, neuroprotection, and transplantation of neuronal pre-cursors. Targeting the root cause, i.e, removing or silencing *mHtt* holds the advantage of ameliorating early events of the disease. Accordingly, approaches that deliver RNAi, miRNAs, or CRISPR/Cas to target the *mHtt* are actively developed (reviewed in Gavgani and García-Domínguez, 2025). Of these, miRNA has proven successful in mice (Sogorb-Gonzalez 2024), and CRISPR/CasRx knock-down of *Htt* was recently reported (Lin *et al*., 2025). Also anti-sense oligos have been developed but faced challenges in clinical trials (Rook and Southwell, 2022). While reducing *mHtt* acts on the early events, clinical strategies need combinatorial approaches with optimized timing to the HD stage. Indeed, mounting evidence shows induced polyQ expression to activate HSF1-driven HSPs expression, and increased HSP levels to protect cells during early phases of polyQ pathology (reviewed in Muchowski and Wacker, 2005). For example, hsf-1 activity and hsp expression ameliorate polyQ stress and extends lifespan in *C.elegans* (Calamini *et al*., 2011; Prahlad and Morimoto, 2011), which has a typical life span of 18-20 days. Growing evidence also shows certain chaperone complexes to maintain aggregation-prone intermediates in a soluble state (Månsson *et al*., 2014; Akaree *et al*., 2025), and balanced co-chaperone networks to be critical in taupathies and HD (Bailus *et al*., 2021; Criaro-Marrero *et al*., 2021; Jeanne *et al*., 2024). Once the polyQ inclusions mature into high-order solid state, however, HSPs likely fail to resolve the aggregation stress and could became trapped into the solid-state complexes. Mice have a typical lifespan of two years, and in the models investigated here, the solid-state aggregates had been formed. Accordingly, we did not find clear *Hsp* induction, but instead, uncovered broadly repressed cellular functions, altered metabolism, and increased autophagy. Hence, work by us and others emphasize the importance of timing and designing the treatment for the disease state; While reducing *mHtt* expression is efficient at early stages, HSP induction could handle soluble intermediates, ideally with tailored co-chaperone network. Once solid-stage aggregates form, management of collapsing cellular processes is required, and neuronal loss could be compensated with transplantation of neuronal precursors. To this end, the highly tissue-specific responses, and disease progression during ageing, challenge treating HD *via* individual proteins or pathways. However, the overarching defects in maintaining cellular processes across tissues raises possibilities for improving energy availability, RNA Polymerase function, lipid synthesis, and autophagy in late-stage HD with solid-state polyQ aggregates. Taken together, our results support stage-timed approaches that reduce *mHtt* expression, induce HSP networks, balance energy needs, improve autophagy, and boost enzymatic de-toxification, in treating HD.

## Materials & Methods

### Sources and alignment of RNA-seq data in *WT*, *R6/2*, *Q175*, and *Hsf1^-/-^* mice

Raw RNA-seq data in *WT*, *R6/2*, *Q175* and *Hsf1^-/-^* mice was obtained from three studies and downloaded as raw fastq files from Gene Expression Omnibus (GEO). The obtained raw data was processed to sequencing depth normalized coverage files, and gene expression counts. The polyA+ RNA-seq data from mouse *quadriceps femoris* under distinct stress conditions was obtained from Neueder *et al*., 2017 (GSE95602). The polyA+ RNA-seq data in 11 tissues of *WT* and *Q175* mice originated from HDinHD consortium (www.HDinHD.org; Langfelder *et al*., 2016; GSE65775), and RNA-seq data in striatum of 2-, 6-, and 12-month-old *WT* and *Q175* mice was from Diaz-Castro *et al*., 2019 (GSE124846). From GSE124845 data, we downloaded the striatal RNA-seq (input). For each sample and replicate, fastq files were obtained with sra-tools (fasterq-dump) and the quality analysed with fastqc (https://www.bioinformatics.babraham.ac.uk/projects/fastqc/). Where needed, the reads were trimmed to remove low-quality ends, and filtered for high quality with fastx-tools (https://github.com/agordon/fastx_toolkit). The reads were mapped to mouse genome annotations (mm10 or GRCm39) with a splice-aware aligners HiSat2 (Kim *et al*., 2019) and STAR (Dobin *et al*., 2013). The obtained .bam files were sorted with samtools (Danecek *et al*., 2021) and converted to bedgraph with bedtools (Quinlan and Hall, 2010), and to bigWig with BedgraphToBigWig (https://www.encodeproject.org/software/bedgraphtobigwig/).

### RNA expression analysis

In each sample, RNA expression was measured with featureCounts (Liao *et al*., 2014) or HTseq (Anders *et al*., 2015), counting alignments to exons for each transcript. The obtained RNA expression was compared between replicates by visualizing log2 converted RNA expressions in xy-graphs and analysing statistical significances with Spearman rank correlation (rho). After ensuring high correlation, replicates were merged into coverage (bigWig) files, and the RNA expression of replicate-merged data visualized with Integrative genomics viewer (IGV, Thorvaldsdóttir *et al*., 2013). Within each IGV browser image, the y-scales of all tracks are identical and linear. Differential gene expression was assessed with DESeq2 (Love *et al*., 2014), which compares variance of expression within replicates to the expression difference between conditions. To call differential expression in comparison of three protein damaging stresses, adjusted p-value < 0.001 and a minimum fold change 1.25 (1.25 for increased, 0.8 for decreased), were required. To account for the higher variation of RNA expression between tissues, adjusted p-value < 0.05 and [log2FC| > 0.5 were used as thresholds. RNAs with less that 10 counts (average of replicates) were considered unexpressed. Differential RNA expression was visualized in MA-plots, showing RNAs with significantly increased expression in red, and significantly decreased expression in light blue. Across the analyses of proteotoxic stresses, heat shock and HSP90 inhibition were matched to their respective controls, HS against NHS, and i90 against iC. In each MA-plot, violin graph, and xy-plot, the exact comparison is indicated. Final RNA expression values are mean expression within replicates against kilobases of transcript length and sequencing depth (FPKM).

### Principal Component Analysis (PCA)

PCA was performed with prcomp R package on log2 transformed RNA expression. In stress response analyses, RNA expression in all genotypes and conditions, including non-heat-shock (NHS), heat shock (HS), vector control (iC), and Hsp90-inhibition (i90) were included. The expression programs were transformed into principal components 1 and 2 that held 33% and 12%, respectively, of the total variance within the all the RNA expression programs. In the xy-graph, each RNA expression program is represented as a dot, and the expression program of unstressed WT mouse (NHS_WT) placed into the origo. The arrows are displacement vectors (mean expression vectors) in the PCA space and show the direction and distance of each RNA expression program, related to the RNA expression program in the NHS_WT mouse. In analysis of RNA expression across tissues, log2 expression means in all *WT* and *Q175* tissues were included. Next, PCA was conducted with brain tissues only.

### Correlation matrices and heatmap

Top 40 most contributing RNAs for variance in principal components 1, 2 and 3 were selected (120 RNAs in total). The expression of the selected RNAs were z-scaled, and the scaled expression visualized on a heatmap, generated with pheatmap package in R. The names of the 120 RNAs are displayed on the heatmap, and the color scale indicates mean (white), high (red, positive z-score), and low (blue, negative z-score) RNA expression.

### Gene Ontology Analyses

Gene ontology enrichments within RNA groups were searched with DAVID tool (Sherman *et al*., 2021) using annotation clustering. The FDR corrected p-values (Benjamini) are shown on the graphs and tables. Due to very small group of GO terms within downregulated genes upon HSP90 inhibition, also kinases are included where uncorrected p-value, but not Benjamini, was significant (indicated with asterisk). Differential gene expression in tissues of *WT* and *Q175* mice was analysed with Gene Set Enrichment Analysis (GSEA; Subramanian *et al*., 2005). Full GO term reports are available in Data S2 (DAVID GO) and Data S5 (GSEA). To identify transcription factors that control genes in the selected groups, we used gene set knowledge discovery with enrichr (Xie *et al*., 2021), with transcription regulatory relationship unraveling TRUSST database (Han *et al*., 2018) with standard settings. Full enrichr reports are available in Data S3.

### Generation of interactive 3D graphs

Stress-induced changes in RNA expression were reported in log2FC as follows. HS: log2(HS_WT / NHS_WT); i90: log2(i90_WT/iC_WT), and R6: log2(NHS_R6/NHS_WT). The log2FC of each RNA was graphed in 3D, one axis per stress using R package plotly (Sievert, 2020), which provides interactive web-based graphics for dynamic rotation, zooming, and point-inspection. The 3D vizualisations were saved as .html compatible files that can be loaded for interactive, user-guided examination by downloading and unzipping File S1 from github.com/Vihervaara/3D-Stress-Cube, and opening the .html file in a web browser. The points are colored based on octant (see schematic cube in Fig. 3). The total distance of each RNA from the origo was counted as sqrt(log2FC_HS^2^ + log2FC_R6^2^ + log2FC_i90^2^). In the interactive visualization, RNAs with a min total distance 1.5 are shown. Data S4 lists all the RNAs and their transcriptional changes during proteotoxic stresses.

### Comparison of transcriptional responses in muscle of chronically stressed mice

After calling significantly induced or repressed genes in each mouse genotype, a unison of induced or repressed genes in a given stress condition was identified. RNA expression changes upon heat shock and HSP90 inhibition was, thereafter, analysed in the unison of induced or repressed genes in the given condition. Violin graphs compare reprogramming of RNA expression in *WT*, *R6/2*, and *Hsf1^-/-^* mice as log2FC. Statistical significance between genotypes is counted with paired student’s t-test, showing the p-value for each comparison on the violin graphs. Since fold change can be high for lowly expressed genes, we also compared differences in RNA induction and expression as total difference, FPKM_treatment - FPKM_control. We first counted the difference in each genotype (e.g. heat shock in WT = HS_WT - NHS_WT), and then compared the total difference between mutant and *WT* as Delta change (e.g. change in *R6/2* - change in *WT*). An example calculation for delta change in heat shock between *R6/2* and *WT* is: (HS_R6 - NHS_R6) - (HS_WT - NHS_WT). The delta change is compared across unisons of induced and repressed RNAs and examples of outlier RNAs are shown in genome browser.

## Author contributions

A.R., I.S., and A.V. conceptualized and designed the study. All authors conducted the computational analysis and interpreted the results. A.R., I.S., and A.V. wrote the manuscript with help from H.L., A.P. and S.A.. A.V. supervised and financed the work.

## Acknowledgements

We thank Jingtian Liu for comments during manuscript preparation. This work was financially supported by Science for Life Laboratory (A.V., SciLifeLab Fellowship), Swedish Research Council (A.V., 2021-02668), and Royal Institute of Technology (A.V.).

