## Supplemental_Figures_S1_to_S7 for "Transcriptional responses to proteotoxic stressors are profoundly diverse and tissue-specific"

**A**

| Mouse | Genotype | Replicate | Treatment | with time | sra download |
|---|---|---|---|---|---|
| 1 | ***WT*** | rep1 | NHS | NHS | SRR5306705 |
| 2 | ***WT*** | rep2 | NHS | NHS | SRR5306706 |
| 3 | ***WT*** | rep1 | HS | HS20min+4hR | SRR5306707 |
| 4 | ***WT*** | rep2 | HS | HS20min+4hR | SRR5306708 |
| 5 | ***WT*** | rep1 | iC | iC_4h | SRR5306717 |
| 6 | ***WT*** | rep2 | iC | iC_4h | SRR5306718 |
| 7 | ***WT*** | rep1 | i90 | i90_4h | SRR5306719 |
| 8 | ***WT*** | rep2 | i90 | i90_4h | SRR5306720 |
| 9 | ***Hsf1-/-*** | rep1 | NHS | NHS | SRR5306697 |
| 10 | ***Hsf1-/-*** | rep2 | NHS | NHS | SRR5306698 |
| 11 | ***Hsf1-/-*** | rep1 | HS | HS20min+4hR | SRR5306699 |
| 12 | ***Hsf1-/-*** | rep2 | HS | HS20min+4hR | SRR5306700 |
| 13 | ***Hsf1-/-*** | rep1 | iC | iC_4h | SRR5306709 |
| 14 | ***Hsf1-/-*** | rep2 | iC | iC_4h | SRR5306710 |
| 15 | ***Hsf1-/-*** | rep1 | i90 | i90_4h | SRR5306711 |
| 16 | ***Hsf1-/-*** | rep2 | i90 | i90_4h | SRR5306712 |
| 17 | ***R6/2*** | rep1 | NHS | NHS | SRR5306701 |
| 18 | ***R6/2*** | rep2 | NHS | NHS | SRR5306702 |
| 19 | ***R6/2*** | rep1 | HS | HS20min+4hR | SRR5306703 |
| 20 | ***R6/2*** | rep2 | HS | HS20min+4hR | SRR5306704 |
| 21 | ***R6/2*** | rep1 | iC | iC_4h | SRR5306713 |
| 22 | ***R6/2*** | rep2 | iC | iC_4h | SRR5306714 |
| 23 | ***R6/2*** | rep1 | i90 | i90_4h | SRR5306715 |
| 24 | ***R6/2*** | rep2 | i90 | i90_4h | SRR5306716 |

**B**

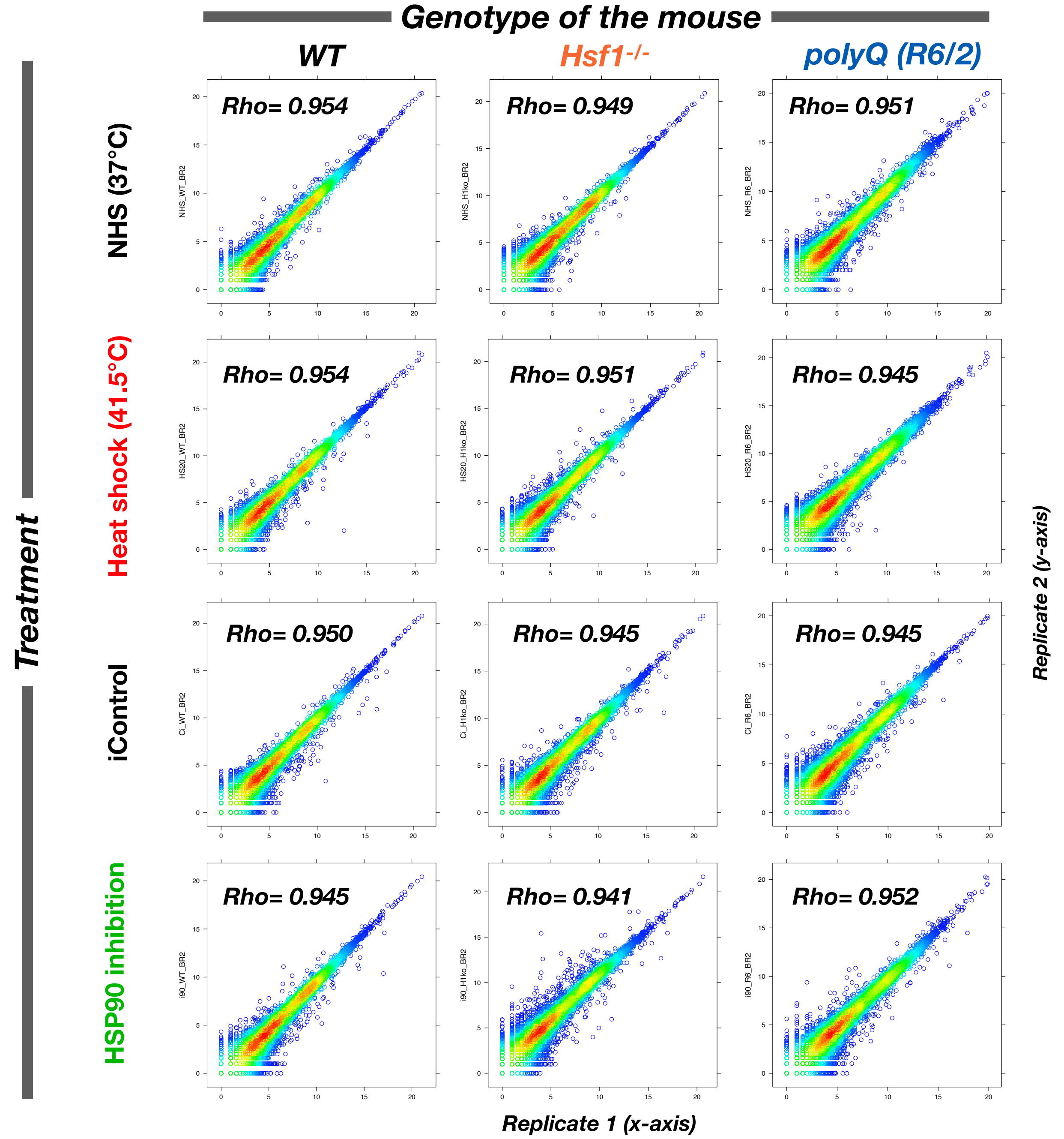


**Supplemental Figure 1. PolyA+ RNA-seq samples and replicate correlation in proteotoxic stress conditions in mouse muscle. A)** Genotype, treatment, and short read archive access code of each sample and replicate used in this study. The data originates from Neueder *et al*., 2017 . **B)** Replicate correlation as log2(counts) on exons of each gene. Spearman rank correlation (Rho) is indicated for each replicate 1 replicate 2 comparison.

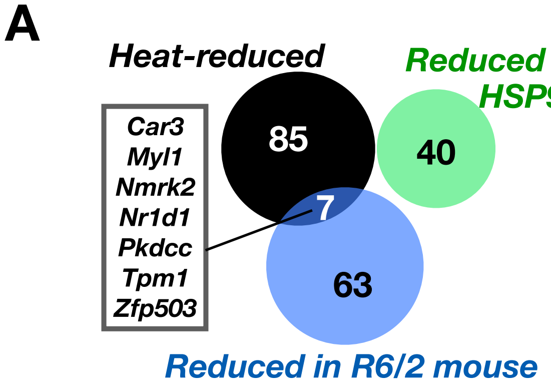


B

| GO terms: heat-repressed | p-value |
|---|---|
| *Myofibril* | *4.5*10-5* |
| *Smooth muscle cell migration* | *2.2*10-2* |

| GO terms: Reduced by HSP90i | p-value |
|---|---|
| *Cytosol* | *1.0*10-3* |
| *Kinase* | *1.7*10-2 ** |

| GO terms: polyQ reduced | p-value |
|---|---|
| *Muscle protein* | *9.9*10-10* |
| *Fatty acid biosynthesis* | *9.3*10-6* |
| *Muscle contraction* | *3.9*10-5* |
| *AMK signalling pathway* | *1.2*10-4* |
| *Lipid biosynthesis* | *1.4*10-4* |
| *Cytoskeleton in muscle cells* | *2.6*10-4* |
| *Myofibril* | *5.7*10-4* |
| *Sarcoplasmic reticulum* | *4.4*10-2* |

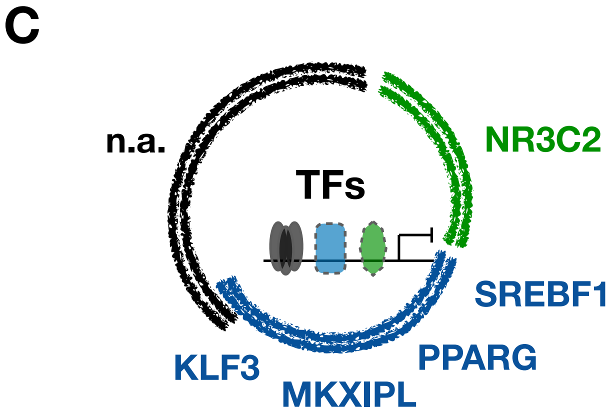


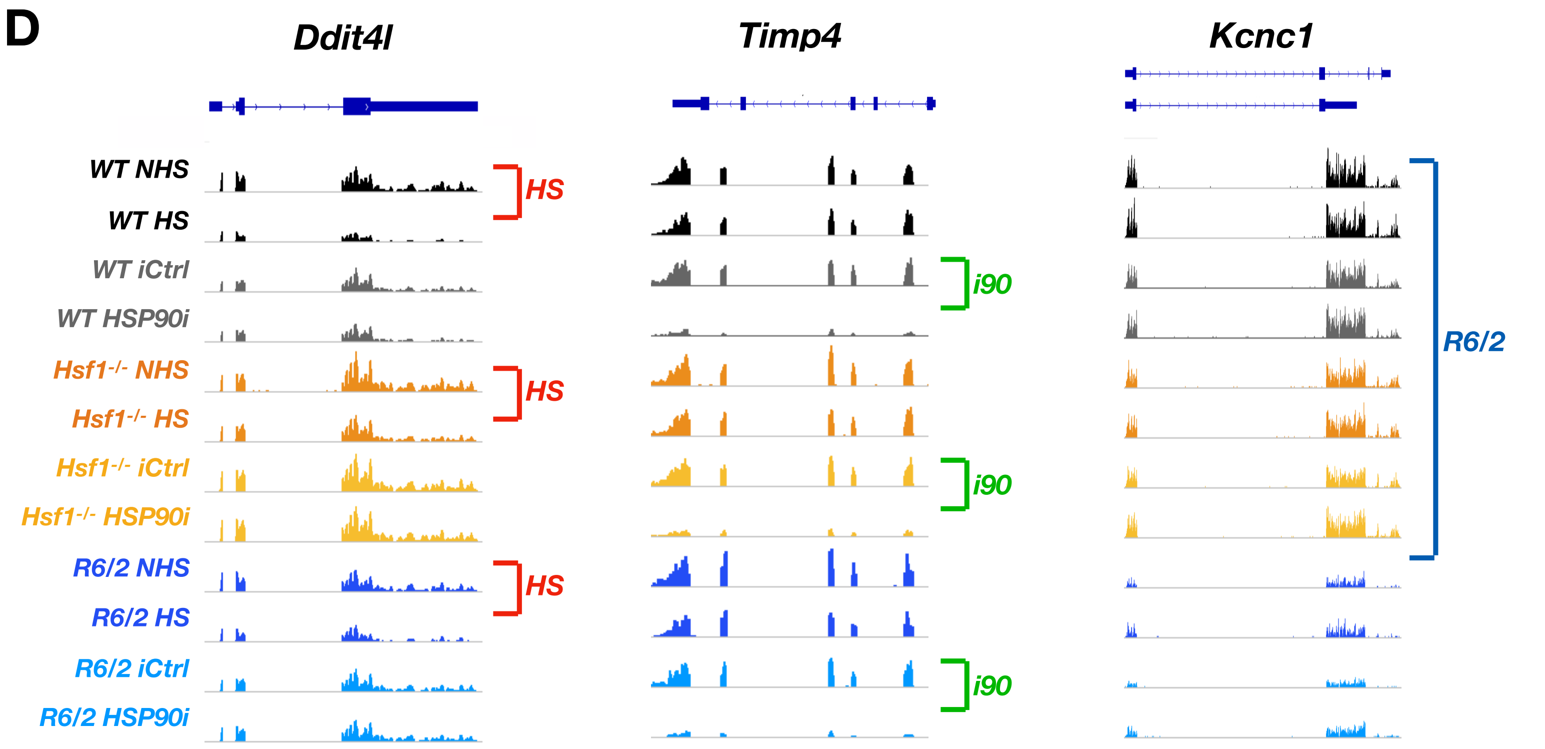


**Supplemental Figure 2. Heat stress, HSP90 inhibition, and polyQ aggregation cause reduced expression of remarkably different RNAs. A)** Significantly decreased RNAs compared in a Venn diagram. Genes that are repressed in two proteotoxic stress conditions are listed in a box. **B)** Enriched GO terms among the stress-repressed genes. FDR corrected p-value (Benjamini) is indicated for each functional category, except for Kinase (HSP90 inhibition), where *non-corrected p-value is given. All categories and their FDR corrected and non-corrected p-values are given in Data S2. **C)** EnrichR analysis of transcription factors predicted to regulate the repressed genes. **D)** Genome-browser examples of genes whose expression is stress-specifically reduced. *Ddit7l* mRNA expression is reduced upon heat shock, *Timp4* upon HSP90 inhibition and *Kcnc1* in *R6/2* mouse.

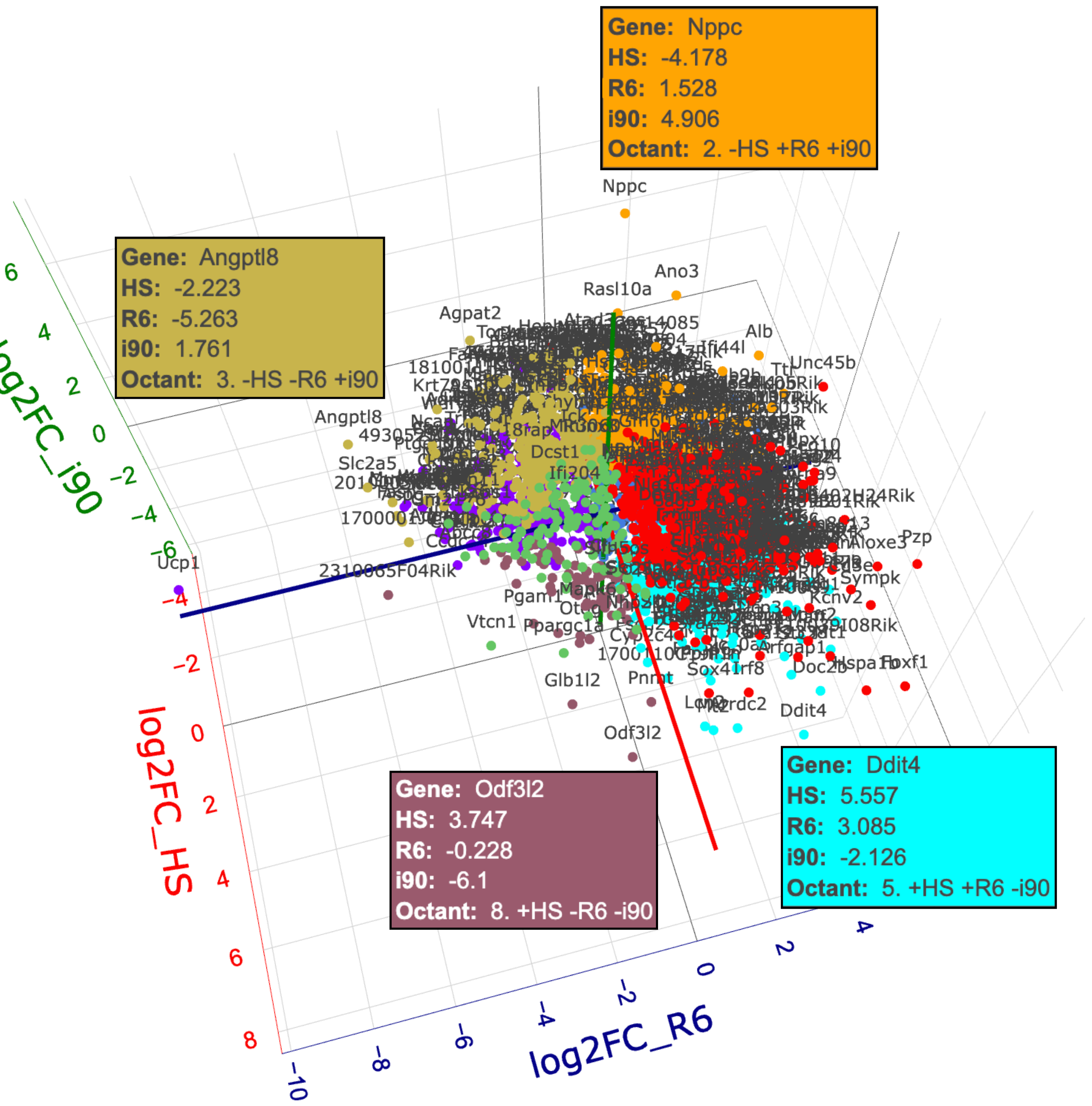


**Supplemental Figure 3. 3D Stress Cube: Interactive visualisation of transcriptional changes upon heat shock, HSP90 inhibition and polyQ aggregation in mouse muscle tissue.** Log2 fold change of example RNAs shown in 3D Stress Cube upon heat shock (HS), HSP90 inhibition (i90), and polyQ aggregation (R6). The RNAs are colored based on the octant, i.e. the direction of the stress-induced changes. *Nppc* (orange), *Angptl8* (gold), *Odf3l2* (burgundy) and *Ddit4* (cyan) RNAs are highlighted from the distinct octants, and the direction (- reduction; + induction) and log2FC in each stress condition shown in the respective infobox. The 3D stress-cube was generated with plotly (Sievert, 2020) and stress-induced changes were counted as follows. Heat shock: log2(HS_WT/NHS_WT). HSP90 inhibition: log2(i90_WT/iC_WT). PolyQ: log2(NHS_R6/NHS_WT). The interactive cube contains RNAs with combined log2FC >1.5, counted as sqrt(log2FC_HS^2 + log2FC_R6^2 + log2FC_i90^2) >1.5. Data S4 lists all transcripts and their respective log2 fold changes.

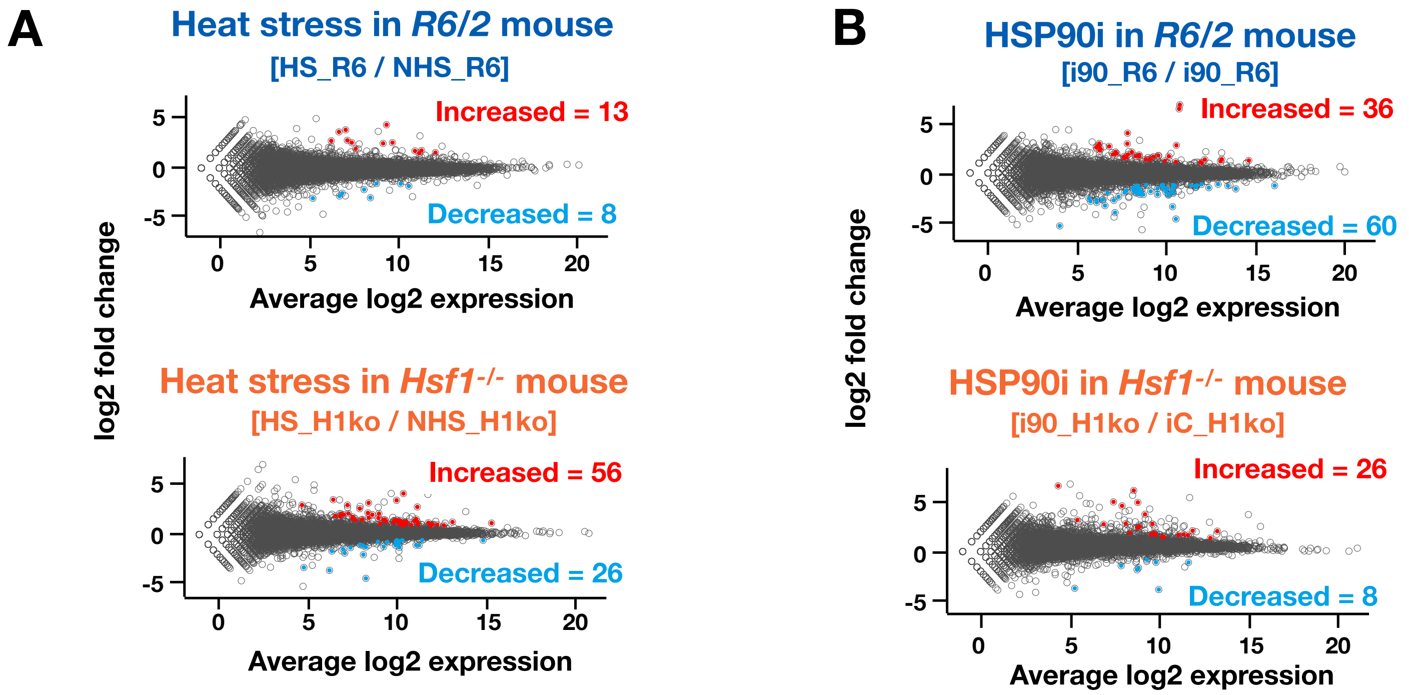


**Supplemental Figure 4. Quantification of stress-induced gene expression in mice under chronic stress. A-B)** MA-plots showing average expression (x-axis) and log2 fold change (y-axis) of RNAs upon heat shock (**A**) and HSP90 inhibition (**B**) in *R6/2* (upper panels) and *Hsf1*$^{-/-}$ mice. Significantly increased (red) or decreased (light blue) were identified with DESeq2 (p-value < 0.001 and minimum fold change > 1.25 for increased and < 0.8 for decreased RNAs).

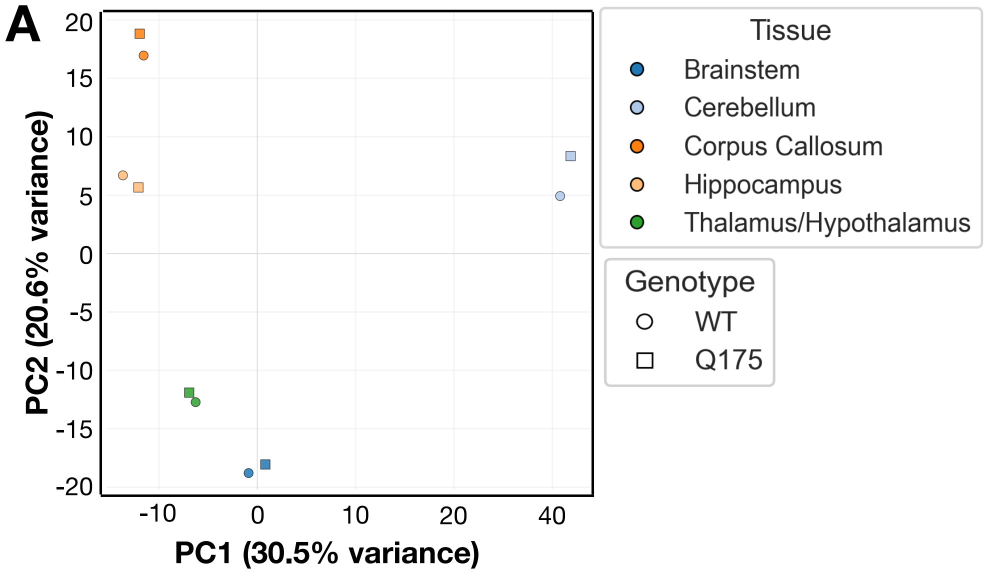


**Supplemental Figure 5. RNA expression in brain tissues of *WT* and *Q175* mice. A)** Principal component analysis of RNA expression programs (log2 expression) in *WT* and *Q175* brain tissues. **B)** Volcano plots showing significantly induced or reduced RNAs (red). To call significance, DESeq2 analysis with p-value < 0.05 and |log2FC| > 0.5 was required. **C)** mRNA expression of *Tmc3* (left panel) and *Abdh1* (right panel) in *WT* (black) and *Q175* mouse brain tissues. Both genes overlap with the end of another, highly expressed gene (*Gm16638* and *Preb1*, respectively).

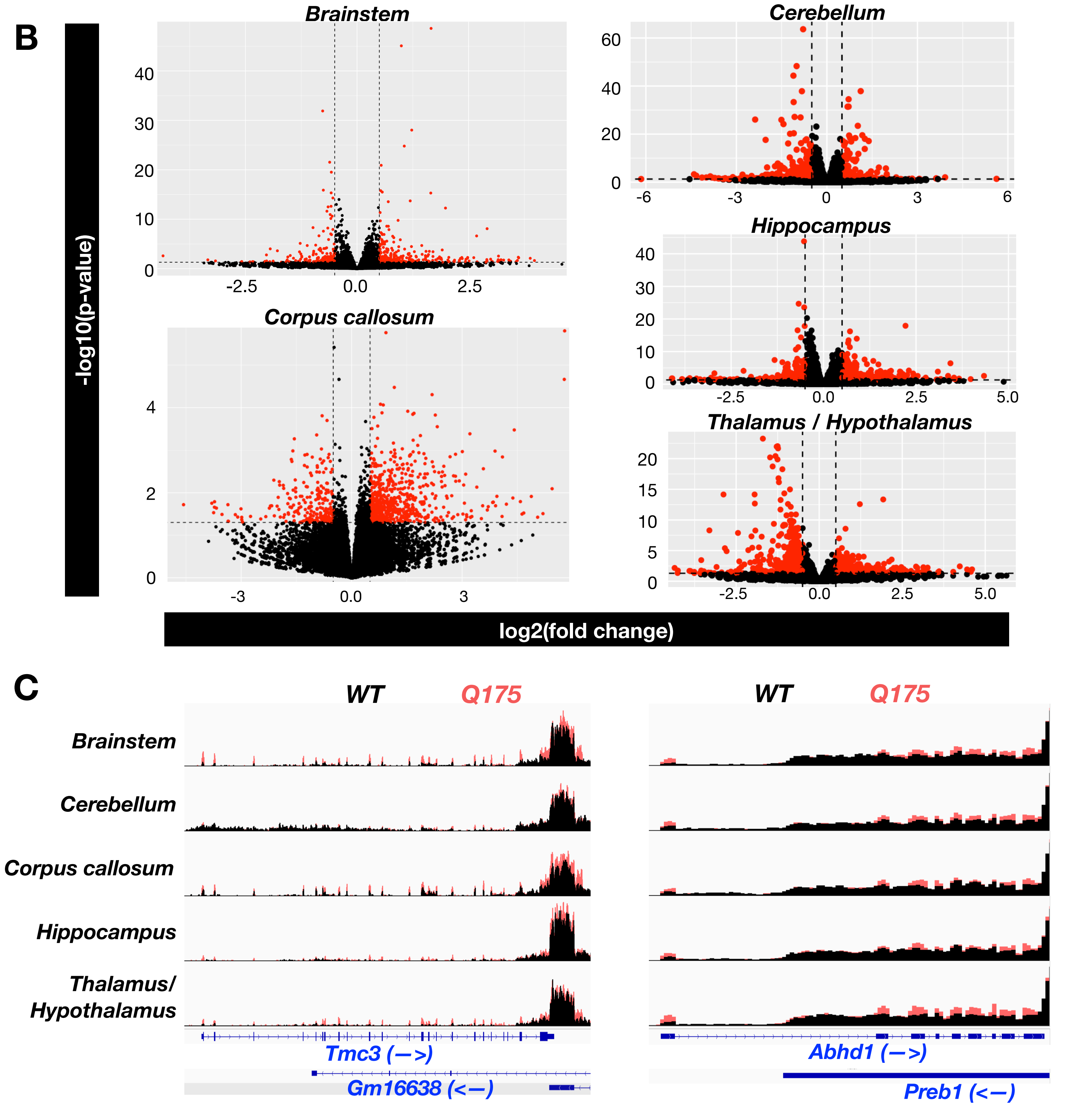

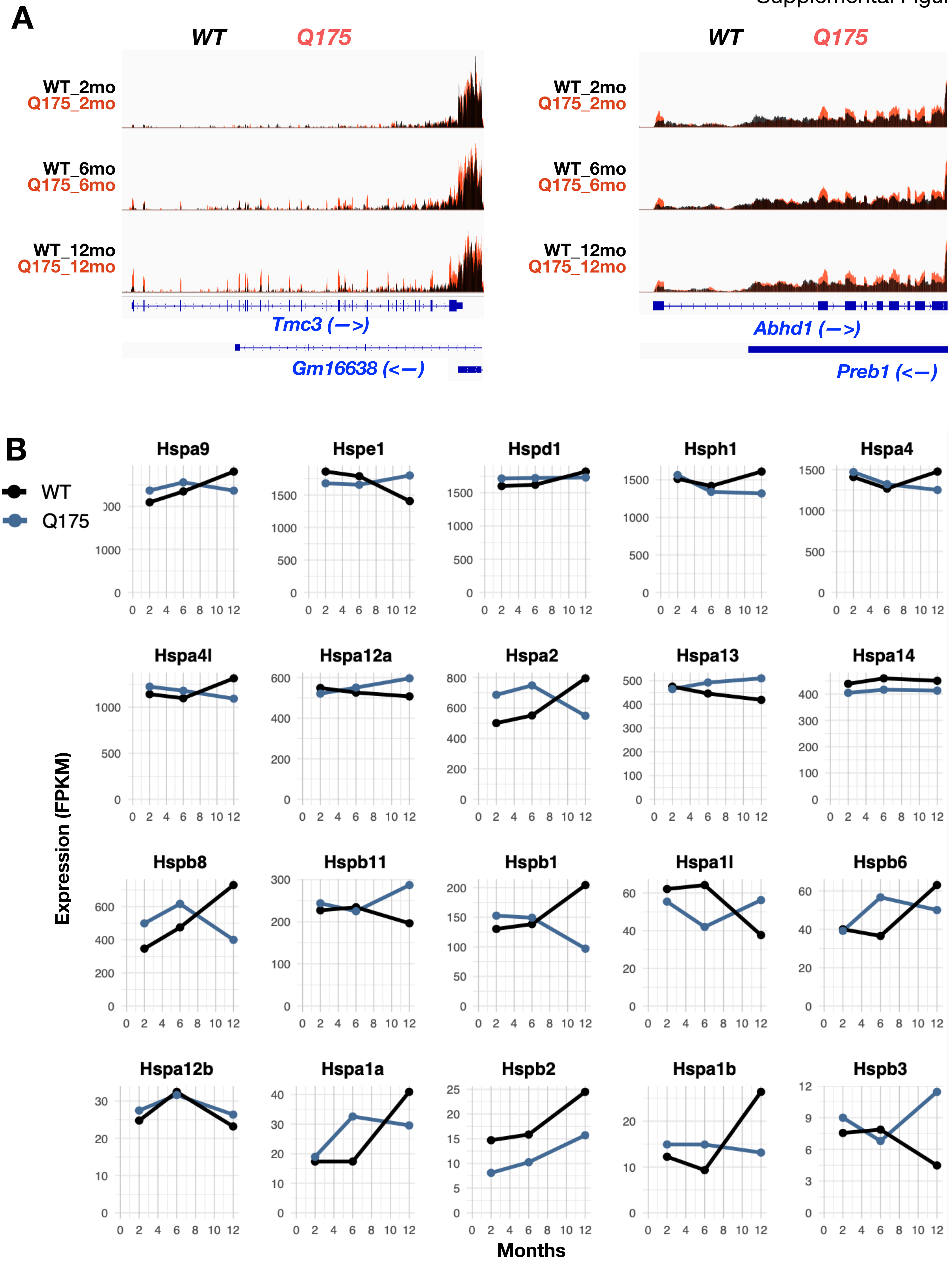


**Supplemental Figure 6. RNA expression in striatum of ageing *WT* and *Q175* mice. A)** mRNA expression of *Tmc3* (left panel) and *Abdh1* (right panel) in *WT* (black) and *Q175* mouse striatum during ageing. Both genes overlap with the end of another, highly expressed gene (*Gm16638* and *Preb1*, respectively). **B)** Expression changes of HSP encoding mRNAs in striatum of *WT* and *Q175* mice during ageing.

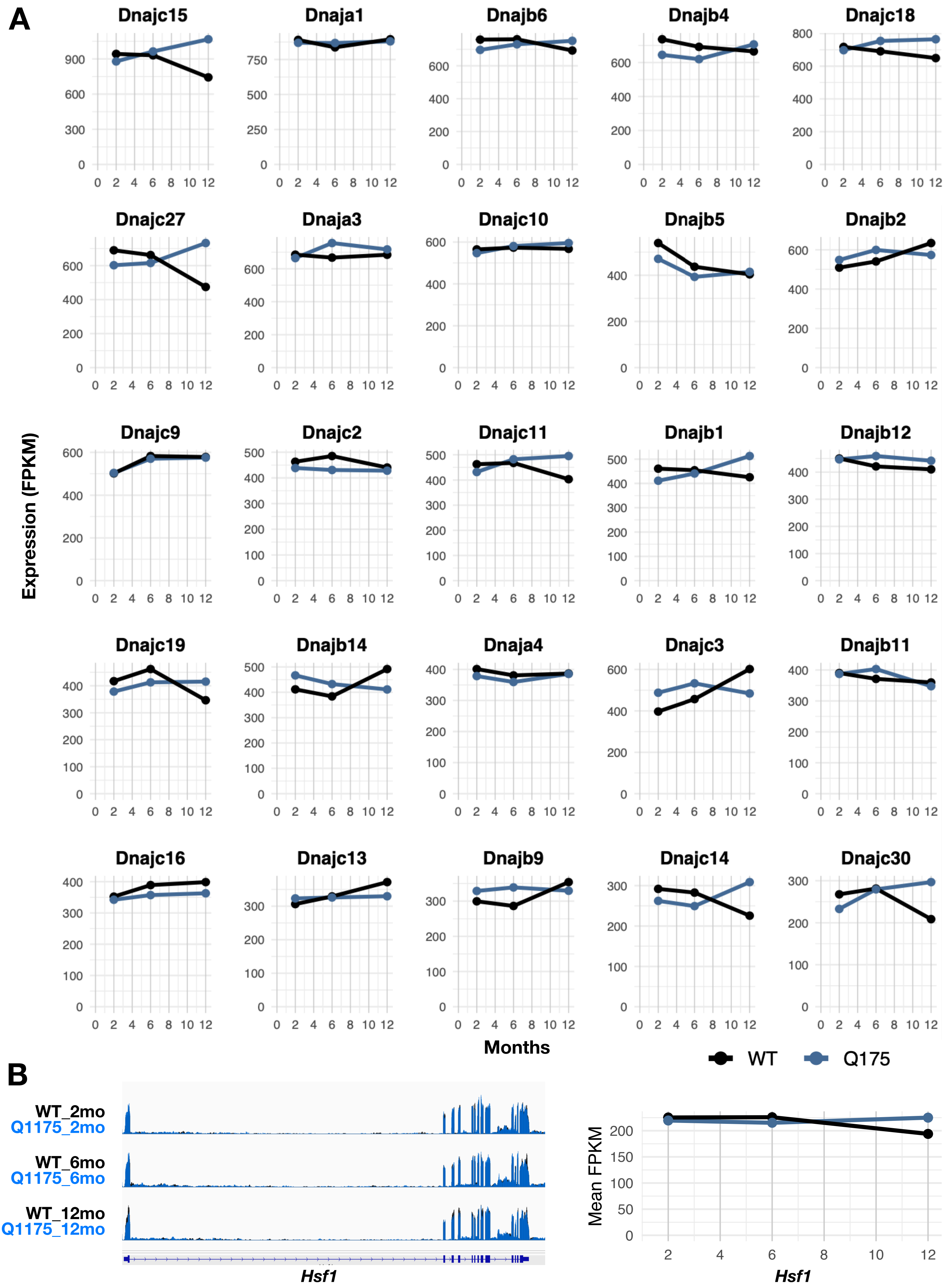


**Supplemental Figure 7. RNA expression in striatum of *WT* and *Q175* mice. A-B)** Expression changes of **A)** DNAJ and **B)** HSF1 encoding mRNAs in striatum of *WT* and *Q175* mice during ageing.
